# Effects of single synonymous substitutions on folding efficiency demonstrate the influence of rare codons and protein structure

**DOI:** 10.1101/2025.03.16.642865

**Authors:** Felipe A.M. Otsuka, Ingemar André

## Abstract

Proteins can misfold during cotranslational folding, but how codon sequences influence this process is not well understood. Here, we develop an *in vivo* assay to comprehensively study the impact of single synonymous substitutions on protein folding efficiency and apply it to the N-terminal domain of E. Coli protein ddlA. We map the influence of codons along the sequence and demonstrate that codons can substantially influence the folding efficiency and that the impact depends on the structure and topology of the protein. A cluster of codons associated with residues in the center of the domain fold strongly influences the folding efficiency. Moreover, substitutions to rarer codons generally lead to increased folding efficiency. A mRNA sequence exclusively made up of rare codons results in higher expression than one with only common codons. Our results highlight the importance of rare codons in cotranslational folding and the relationship between codon sequence and protein structure.

## Introduction

Proteins are synthesized during translation by decoding nucleotide triplets (codons) in mRNA, which directs the ribosome to add amino acids to a growing polypeptide chain. While it is well established that the amino acid sequence contains the blueprint for the native structure of proteins, it is less understood how the folding of the nascent chain is affected by the sequential nature of the translation process (1). The cotranslational folding of the peptide chain as it emerges from the ribosome can help guide the protein toward the correct final structure (2) but can also lead to misfolding (3). This misfolding may not always be resolved with spontaneous refolding and can result in a protein with compromised function, cell stress responses, and aggregation (4).

Folding during translation is not only affected by the amino acid sequence but also by how the amino acids are encoded as codons (5, 6). The genetic code is degenerate, with 18 of the 20 standard amino acids encoded by multiple synonymous codons. Individual codons can significantly influence translation speed, as the time required for a cognate tRNA to deliver an amino acid to the ribosome varies between codons. When a codon exhibits longer decoding times compared to its synonymous counterparts, it is classified as "rare" (7). mRNA regions enriched in rare codons are associated with ribosome pausing, suggesting a coordinated mechanism in which codon sequence within a gene regulates ribosomal speed (8). However, the precise impact of codon-dependent translation kinetics on protein folding remains elusive, as it is still unclear which parts of the linear amino acid sequence are most sensitive to changes in translation speed.

The role of codon usage in modulating folding has been described as a secondary layer of genetic encoding within the standard amino acid sequence (9). Evidence suggests that specific sets of codons can dictate distinct folding pathways during translation to guide the folding of proteins toward their correct native structures (10, 11). Data also indicate that even subtle codon changes within the coding sequence (CDS) can significantly alter the folding trajectory of a nascent protein. Single synonymous mutations—or a pair of them—within the CDS have also been shown to modulate protein function (12–14). Interestingly, both “slow” and “fast” decoded codons could help prevent misfolding. Slow codons enable more time to find the native contacts, while fast codons can upregulate speed in regions prone to misfolding (15, 16).

Bioinformatic studies have shown that regions of the mRNA encoding secondary structure of proteins contain conserved patterns of optimal and nonoptimal codons in some regions of genes, suggesting that codons are often under selective pressure (17, 18). The influence of codon choices on overall cell fitness (14, 19) has been investigated, and there is ample data linking non-optimal codon choices to poor protein expression, misfolding, and aggregation (3, 20, 21). Nonetheless, there is a shortage of experimental data providing the basis for a mechanistic model of how synonymous codon choices can prevent misfolding. This type of data is necessary to answer some critical questions remaining on the influence of codon choices on cotranslational folding: which codons within a CDS influence cotranslational folding the most? How many codon changes, in general, are required to see a substantial effect on folding? How important is the protein structural environment associated with a codon in explaining its influence on folding?

To increase our understanding of the impact of individual codon choices on folding it would be valuable to have a robust and general experimental method to couple folding efficiency and codon sequence. This would also provide an alternative source of data to improve models of codon selection for optimization of protein production in biotechnology. In this study, we systematically assess the impact of single-codon substitutions on protein misfolding using an *in vivo* protein folding assay. This work extends a previously developed system (22) for measurement and comparison of the folding efficiency of non-synonymous mutations. We have demonstrated how this system can be used to study the correlation between protein stability and heat shock response in E. Coli (23). This folding assay can be used for high-throughput stability measurements and accurate identification of stabilizing mutations in a protein (23). Previous studies correlating codon sequence and functional expression have been largely restricted to fluorescent or bioluminescent proteins due to their convenient real-time detection (24–26), by library-based selection of synonymous variants. While these studies have yielded significant insights, they have limitations in terms of diversity in protein structure and function. There is also a lack of systematic investigations of the impact of single codons on folding.

To select a protein sequence to explore with our system to measure folding efficiency, we reasoned that it would be beneficial to use a native E. coli protein classified as non-essential, thereby minimizing potential biases associated with codon usage constraints. Furthermore, the protein should exhibit limited refolding ability upon denaturation, even in the presence of chaperones (27). This enables the measurement of translation effects on folding with minimal interference of the refolding reactions in the cytosol. Based on these criteria, we selected D-alanine—D-alanine ligase A (ddlA) (28) as a model protein. ddlA is predicted to be a client target of the chaperone Dnak (29) (fig. S8), making it more sensitive to protein quality control-mediated stress responses. Additionally, ddlA exhibits thermal stability (Tₘ = 61°C) above the average for proteins in E. coli, while its protein synthesis rate ranks among the lowest (30). We carried out a deep synonymous scan in the N-terminal domain of ddlA and demonstrated that single mutations can have a substantial influence on folding efficiency. Counterintuitively, our results show that changes from common to rare codons consistently enhance the folding efficiency of this protein. An extremely biased sequence, with only rare codons, expresses and folds much more efficiently than a sequence with common codons. Furthermore, by mapping the effect of single codons onto the three-dimensional structure of ddlA, we can show which residue contacts are influenced by the effects of codon on folding.

This work offers a detailed view of how the influence of codons varies with sequence location and structure using a general methodology that can be applied broadly to different proteins. Such investigations can lay the groundwork for a generalized framework to understand the impact of codon choices on cotranslational folding.

## Results

### Measuring folding efficiency of single synonymous mutants of ddlA

In this study, we sought to systematically investigate how synonymous mutations impact the folding efficiency of a protein sequence expressed in E. Coli. Building on a method developed to characterize the stability of protein variants *in vivo*, we built a sensor system (Fig.1a, b, and fig. S1) to measure the folding efficiency of synonymous mutants. The abundance of the gene of interest, ddlA, is measured through a fusion with a red fluorescent protein (ddlA-RFP). To more closely control the level of protein expression, we replaced the T7 promotor in the original stability sensor with an araBAD promoter placed upstream of the coding region (Fig. 1a, and fig. S1). The presence of unfolded/misfolded protein in the cytosol induces a heat shock response through the sigma factor 32 (σ^32^), which triggers the protein quality control system (PQS) (22). σ^32^ is a transcription factor usually bound to inactivate chaperones, *e.g*., DnaK, but when misfolded proteins accumulate (31, 32), chaperones lose σ^32^ to assist the fold of misfolded proteins. The second part of the folding sensor is a superfolder green fluorescent protein (GFP) (33) (Fig. 1b) under the control of a DnaK promotor from E. Coli. We have previously shown that the ratio of red to green color in this sensor system is an accurate folding sensor in the cell and demonstrated that it can be used to identify stabilizing mutations in proteins (22, 23). In this study, we use this approach to investigate the impact of single synonymous codons on folding efficiency.

**Figure 1.**
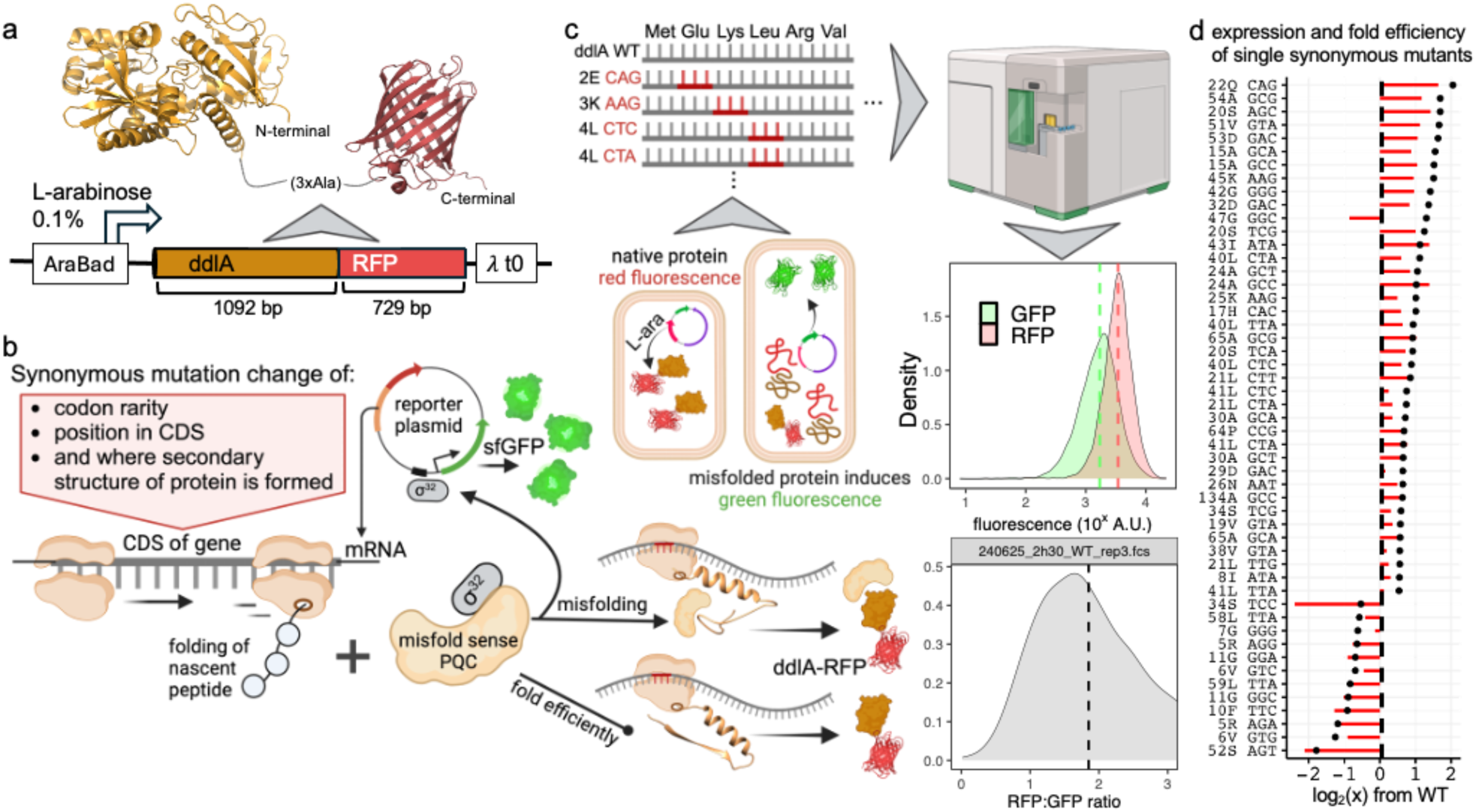
Single synonymous mutants in the ddlA CDS affect folding efficiency, as measured by flow cytometry. **a**) ddlA-RFP cloned in the reporter plasmid. **b**) Codon changes in the CDS may induce cotranslational misfold states which are detected by the PQS, leading to σ^32^ release and activation of GFP transcription via the reporter plasmid. Conversely, when ddlA folds efficiently, PQS activation is reduced and less GFP is expressed. **c**) Overall workflow of the assay: depiction of the reporter plasmid outcomes inside the cell. Each codon variant from the parent plasmid ddlA WT is individually transformed in E. coli and analyzed by flow cytometry. The fluorescence intensities and RFP/GFP ratios for each event are recorded. Dashed lines represent median fluorescence values, which are used in downstream analyses. **d)** FC ratios > 0.5 or < −0.5 of each mutant (black dot) and ddlA-RFP levels (red bars) compared to the ddlA WT. All values are shown in **fig. S7**.

After induction of protein expression, cells are analyzed by flow cytometry, which allows for measurement of red and green fluorescence in individual cells in the expressing population (Fig. 1c). The fold efficiency can then be estimated as the ratio ddlA-RFP and GFP fluorescence signal, reflecting the degree of misfolding normalized by the ddlA-RFP WT expression (Fig. 1b – d, Fig. S2). The araBAD promoter provides a key advantage in tightly regulating ddlA-RFP expression, effectively minimizing background expression prior to induction (Fig. S3). Another benefit of the araBAD system is that the expression rate of the gene-of-interest can be titrated, reducing the risk of saturation of the PQS and increasing the dynamic range of the folding sensor. Compared to expression with araBAD (0.2% L-arabinose), the T7 promoter results in a lower abundance of ddlA- RFP yet a higher chaperone response, potentially indicating some saturation of the PQS (Fig. S3). 0.2% L-arabinose is the standard concentration for araBAD induction. To reduce the risk of saturation a concentration of 0.1% was used, while still giving rise to a good fluorescence signal. The benefits of araBAD versus T7 is further enhanced in the 96-deep well format used in this study (fig. S5), where reduced oxygenation and a delayed expression profile may leads to more leaky expression with T7 (fig. S3).

We scanned for single codon substitution in the N-terminal domain of ddlA from codon 2 to 66 and 123 to 142, a limit of 213 possible single variants to analyze. A total of 93 single variants were confirmed and compared by taking the average over the median RFP/GFP fluorescent ratio for 3 replicates, normalizing with the value for the wild-type (WT) codon sequence (fig. S7). In Fig. 1d the relative fluorescence ratios of all synonymous mutations characterized in this study is shown (more details below), with single changes resulting in up to four times changes in RFP/GFP ratios compared to WT (Fig. 1d, fig. S2). A positive fold-change (FC) calculated as log2(ratio Mut/ratioWT) indicates greater folding efficiency compared to WT, while negative values indicate lower efficiency. These results shows that single codon changes can cause substantial effect on protein folding (FC >> 1 or FC << −1) but also indicate that most mutations with substantial impact lead to increased folding efficiency.

### Position-specific and effects of codon rarity on folding efficiency

To understand how folding efficiency is affected by codon choices with respect to position, secondary structure, and topology a deep synonymous scan was carried out in a segment of the protein. We focused on the beginning and end of the N-terminal domain of ddlA, which is the first to emerge from the ribosome tunnel, corresponding to amino acids 2 to 66 and 126 to 142 (fig. S6). Fig. 2a shows the normalized RFP/GFP ratio of synonymous mutants relative to the WT ddlA and a mapping of the average effect in folding efficiency onto the predicted structure of the N-terminal domain. The correlation between RFP/GFP ratio and RFP (Fig. 1d) is higher than with GFP (Fig. 2b). The mean values of all the alternative synonymous codons at each position are in fig. S7. Significant changes in the protein fold efficiency were detected at three structured regions of ddlA: i) from ^5^Arg to ^11^Gly. Here, the impact was likely due to decreases in ddlA-RFP expression (Fig 2a, Fig. 3c, and fig. S10). Which can be attributed to changes in stability of mRNA at 5’ end, hampering translation initiation (26); ii) from ^21^Ser to ^26^Asn where the highest RFP/GFP ratios were detected; and iii) from ^42^Gly to ^54^Ala which concludes the initial structural motif of the N-terminal domain of ddlA (Fig. 2a, bottom). The codon with the largest positive influence on the folding efficiency is found for ^22^Gln-CAG in the first half of the first α-helix of the protein (Fig. 2a-c), suggesting that the wildtype codon choice could increase the likelihood of misfolding, possibly the helix itself. As the chain becomes longer it can emerge outside the exit tunnel of the ribosome (2) (fig. S8). This is predicted to happen at the end of the third beta-sheet (^42^Gly to ^54^Ala). To further validate the results, we selected a few variants (^24^Ala-GCC, ^28^Val-GTG and ^38^Val-GTG) and followed the kinetics of expression after induction over a time span of 1.5-4.5h (fig. S9 and S13). Overall, the results show that the average impact of codons on fold efficiency is substantial at the beginning of ddlA, while the effects on the last segment scanned, residue 126-142, are mostly neutral, except a variant ^134^Ala (GCT_WT_ to GCC_mut_) which has a positive effect on folding efficiency (Fig. 3c) and is located in the first half of an α-helix. Noteworthy is also that beyond the very beginning of the sequence, folding efficiency can be increased from WT by selecting alternative codons.

**Figure 2.**
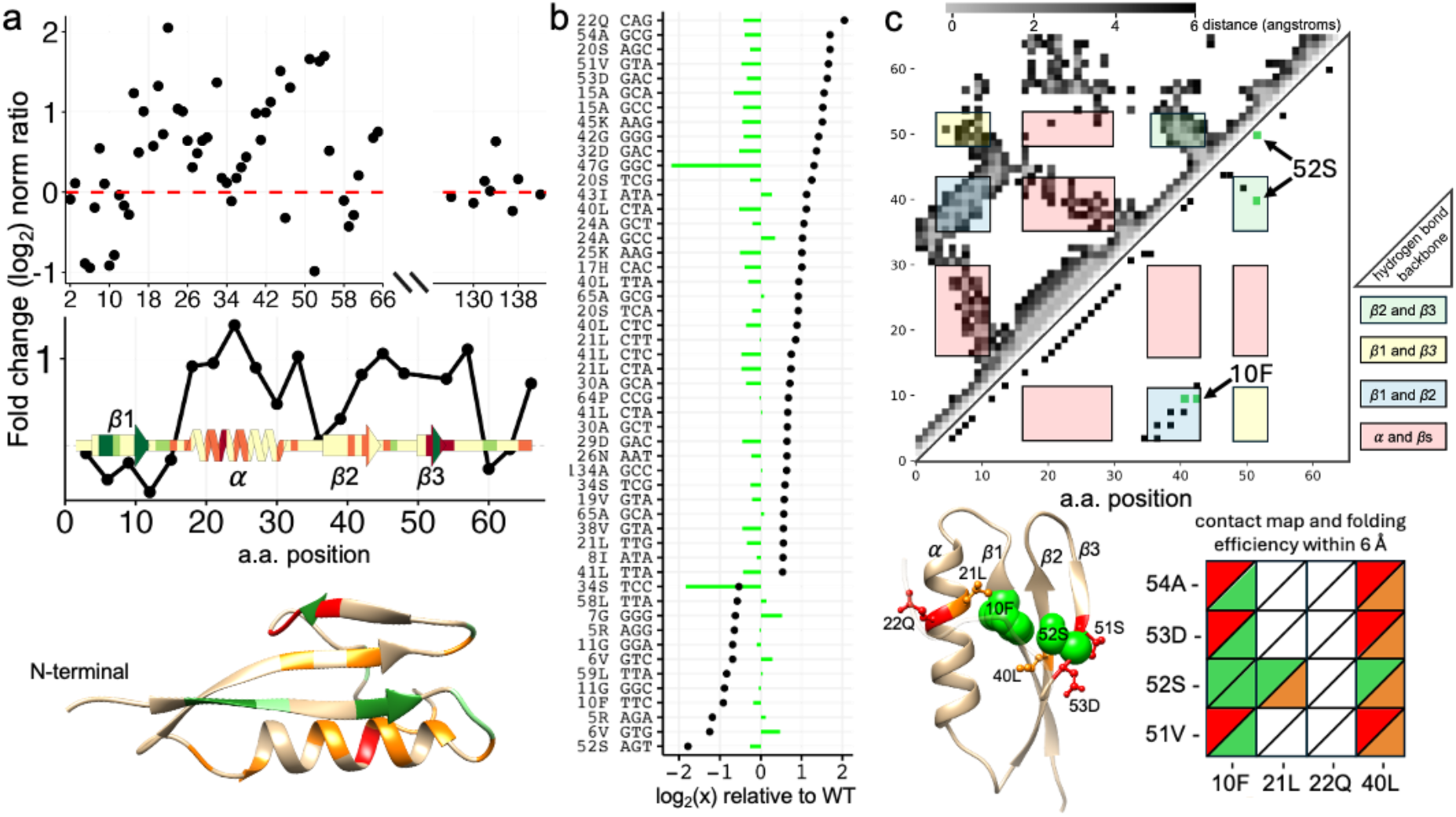
Mapping fold efficiency data of single synonymous mutants onto the ddlA structure. **a)** Average FC of RFP/GFP ratio of synonymous mutants per position relative to WT ddlA. Plot at the bottom depicts three residues binned from position 1 to 66, overlaid with the predicted protein secondary structure (34) of ddlA colored as shades of red for substantial ratios higher than the WT and shades of green for ratios lower than the WT. Bellow the plot, the AlphaFold2 predicted tertiary structure (35) is shown using the same color scheme. **b)** Mean of FC ratios > 0.5 or < −0.5 of each mutant (black dot) and GFP levels (green bars) compared to the ddlA WT. **c)** Contact map of the center of mass for residues 1 to 66 of ddlA, showing atomic proximities within 6 Å (upper triangle), while the lower triangle shows hydrogen bonds from backbone interactions. The corresponding tertiary structure (bottom left) marks residues exhibiting the most substantial changes in the RFP/GFP ratio upon codon substitution. The bottom-right matrix depicts the presence (colored with the same scheme in “a”) or absence (white) of contacts between specific residues.

**Figure 3.**
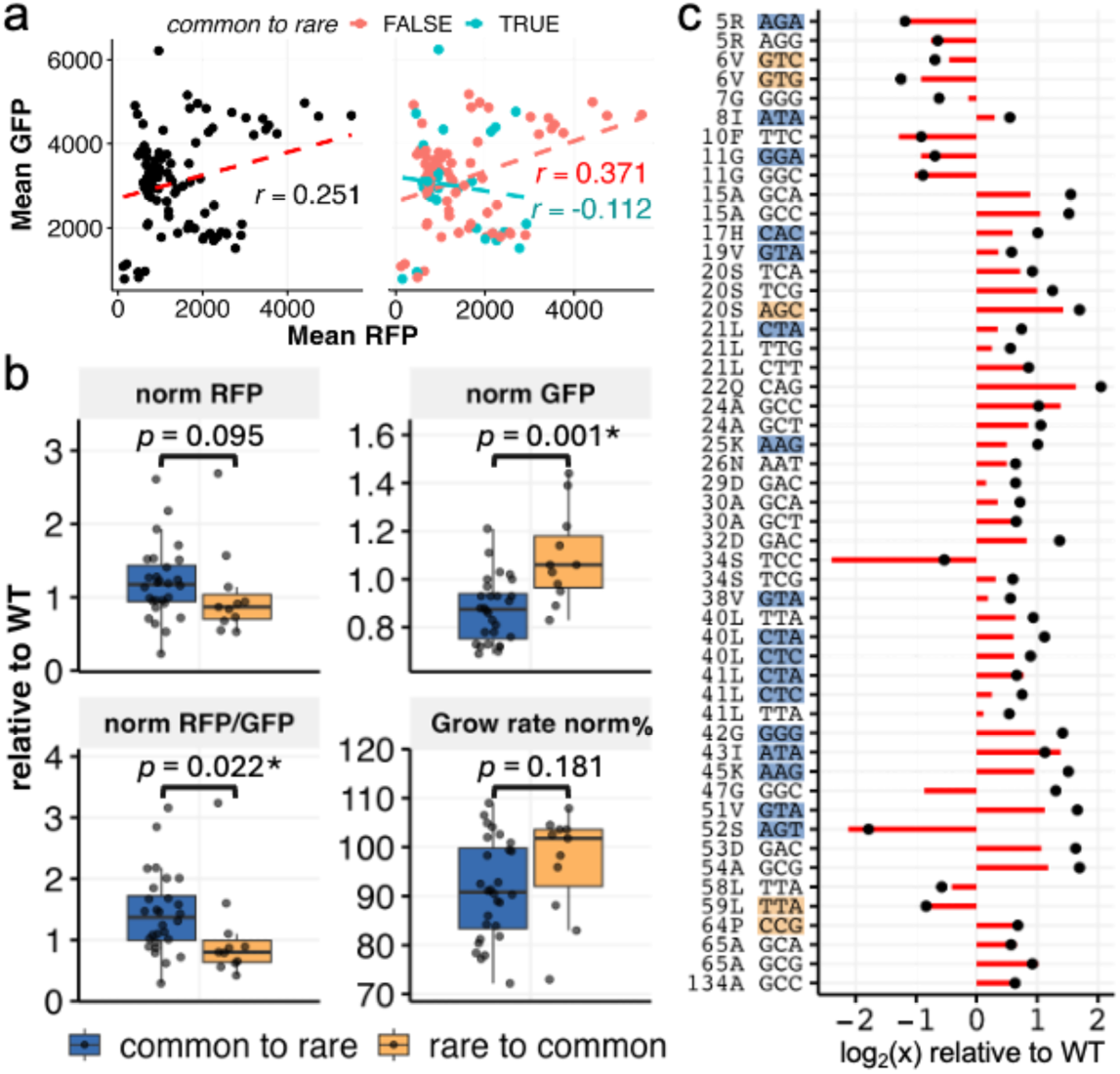
Effects of codon rarity changes in single synonymous mutants of the ddlA CDS. **a)** Scatter plots are the Pearson correlation of mean fluorescence measurements of all single synonymous mutants (left) and after isolate by change in codon rarity (right, fig. S11, and DataS1). **b)** Analysis by category of codon change in the WT ddlA CDS (DataS1). Box plots are normalized values relative to WT and comparisons were tested with Wilcoxon rank-sum test. **c)** FC ratios > 0.5 or < −0.5 of each mutant (black dot) and RFP levels (red bars) compared to the ddlA WT (all data in fig. S7) with change in codon rarity highlighted by the same color palette in as “b”.

To investigate how the effect of codon choices on folding efficiency correlates with the topology and tertiary interactions in the N-terminal domain we mapped folding efficiency values onto a contact map (Fig. 2c). Such analysis can highlight residues where synonymous mutations had the most significant impact on folding efficiency. The analysis reveals a close contact between residues ^10^F and ^52^S, positioned within β-strands β1 and β3, respectively. Both exhibit comparable and negative effects on folding efficiency, suggesting that synonymous mutations at ^10^Phe (TTC) and ^52^Ser (AGT) may contribute to a shared misfolding pathway that disrupts the packing of this β- sheet motif. Interestingly, as translation progresses, codons flanking ^52^S exert a pronounced positive influence on folding, suggesting a codon-specific compensatory mechanism in this region, where the nascent protein undergoes cotranslational folding, either within or outside the ribosomal exit tunnel (2) (fig. S8). This finding underscores the intricate role of unique codon choices in orchestrating cotranslational folding within this structurally sensitive β-sheet segment. Notably, a set of connected residues emerges within this N-terminal motif (Fig. 2c, bottom left), linking residues where synonymous mutations exert significant effects on folding, specifically ^22^Q (CAG), ^21^L (CTA, TTG, CTT), ^10^F (TTC), and ^52^S (AGT). This pattern suggests that these residues play a critical role in the folding of the N-terminal domain and highlights the importance of their corresponding codons in modulating the folding landscape of ddlA. Interestingly, the rather unconventional β-sheet architecture, consisting of two parallel and one antiparallel β-strand, may play a role in the increased complexity of folding within this motif. This topological complexity could explain why codon usage plays a pivotal role in fine-tuning the folding process, rendering this region particularly sensitive to translation rates.

Previous work has established the relationship between protein expression rate and the relative concentrations of cognate tRNA, with rare codons typically associated with lower protein translation rates (5). We categorized substitutions into two categories, common to rare and rare to common, and analyzed their properties. In the overall data, there is only a weak correlation between ddlA-RFP abundance and chaperone response (Fig. 3a). However, when the substitutions are analyzed in the two categories, substitutions to rare codons have an opposite correlation compared to substitutions to common codons (Fig. 3a and fig. S11). Moreover, comparison of opposite codon changes shows that exchange from common codons to rare codons significantly contributes to fold efficiency due to lower expression of GFP (Wilcoxon ranked test, *p* = 0.001), while the change from rare to common is detrimental to fold efficiency (Fig. 3b and fig. S12, S13). Grow rate was not significantly different between the cells analyzed in the codon rarity change (Fig. 3b, and fig. S13), further supporting a direct relationship between codon rarity and protein folding. This result highlights the importance of rare codons in the folding of ddlA while common codons are associated with misfolding during cotranslational folding.

### Rare codon frequency in ddlA CDS does not hamper expression, but abundant codons does

After finding that rare codons favor folding efficiency more than common codons in ddlA, we decided to compare the variants where the entire coding sequence is made up of common or rare codons (Fig. 4a, b) based on the codon usage of E. coli (36). One variant is made of rare codons at each position and has a very low Codon Adaption Index (CAI) of 0.351. A second variant was synthesized with only common codons with a CAI of 1.0, for comparison, the WT ddlA has a CAI of 0.722. The expression kinetics of both variants were determined in parallel with the WT ddlA- RFP (Fig. 4c). Surprisingly, we saw the CAI 0.3 sequence actually express as well as the WT sequence while the expression of the CAI 1 variant was considerably reduced. In a second experiment with longer expression times to allow for both codon variants to reach comparable levels of ddlA-RFP signal, the fold efficiency as measured by the RFP/GFP ratio of the CAI 0.3 variant was still better if compared by the CAI 1 in two different post-induction times (4h and 6h, Fig. 4d, and fig. S14).

**Figure 4.**
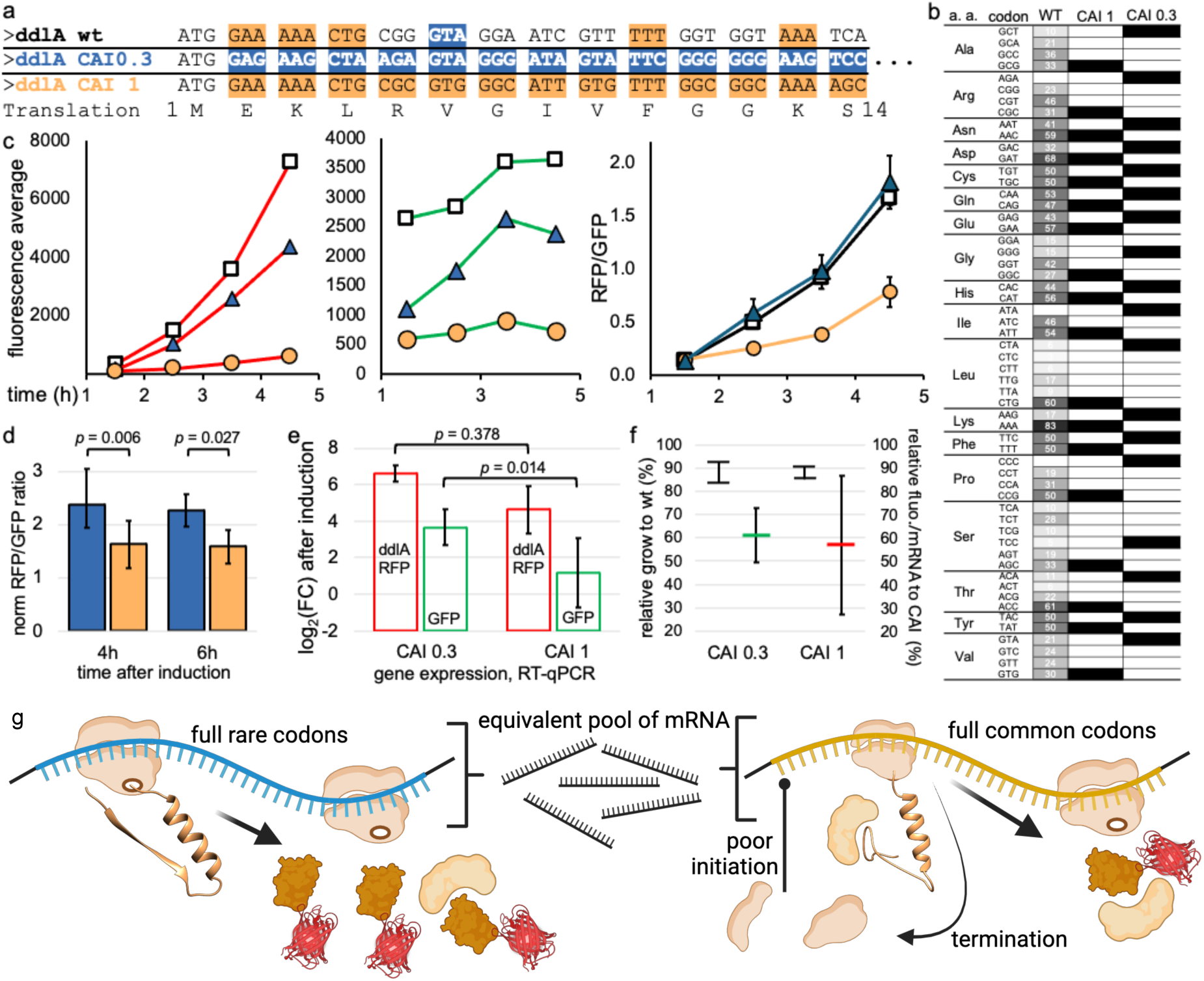
Sequence analysis, expression, and RT-qPCR of single codon frequency per amino acid. **a, b)** Partial view of the codon sequence (positions 1–14) displaying one type of codon per amino acid, along with a frequency map of codon usage in WT ddlA and CAI variants with single codon frequency. **c)** Kinetics of ddlA-RFP expression in Erlenmeyer flasks (20 mL cultures) for the WT CDS (squares), CAI 0.3 (blue triangles), and CAI 1 (yellow circles) showing the mean readouts for RFP (left), GFP (middle) and RFP/GFP ratios (right) of three biological replicates. **d)** Normalized RFP/GFP ratio for CAI 0.3 (blue) and CAI 1 (yellow) induced in a 96-deep well plate at two different time points with biological triplicates. **e)** Fold change in ddlA-RFP and GFP transcripts levels relative to non-induced cells. **f)** Left axis: relative growth of CAI 0.3 and CAI 1 compared to WT during induction in a 96-deep well plate (error bars indicate standard deviation). Right axis: colored stripes represent the percentage of GFP/transcript (green) in CAI 0.3 compared to CAI 1, and the RFP/transcript (red) in CAI 1 compared to CAI 0.3. Data from RT-qPCR represent the mean of two biological and three technical replicates (fig. S15). **g)** Schematic model illustrating the consequences of both CAI variants in the CDS of ddlA.

These results suggest that more well-folded protein is produced with rare codons, indicating that cotranslational folding can result in misfolded states with faster codons. Nonetheless, it is feasible that part of this effect is due to changes in mRNA abundance rather than protein translation and folding. To investigate this, we used RT-qPCR to quantify the mRNA transcripts of ddlA-RFP and GFP from the same experiment (Fig. 4d) and in relation to non-induced cells. The mean of ddlA-RFP transcript levels in the CAI 0.3 sequence is slightly higher than for the CAI 1 sequence (Fig. 4e, fig. S15), but this difference is not significant (*p* = 0.378 two tail t-test with two biological replicates). This suggests that the higher levels of ddlA-RFP in CAI 0.3 compared to CAI 1 are from post-transcription events and cannot be explained by the number of copies of mRNA.

Furthermore, we also saw the mRNA levels of GFP were significantly higher with the induction of ddlA-RFP with CAI 0.3 sequence compared to the CAI 1 (*p* = 0.014 two tail t-test, Fig. 4e, and fig. S15). However, GFP abundance per transcript in CAI 0.3 corresponds to 60% of the value for in CAI 1 (Fig. 4f, green stripe), indicating that the CAI 1 sequence gives about 40% more GFP signal per unit of transcript. The ddlA-RFP fluorescence per transcript of the CAI 0.3 appears to be 40% higher than in CAI 1 (Fig. 3f, red stripe), indicating that CAI 0.3 makes more protein per transcript, but the variability of measurements for ddlA-RFP transcript in CAI 1 is too big for any quantitative conclusions. Differences in protein levels per transcript can be explained by mRNA stability and degradation (37). Taken together, we conclude that mRNA copies of ddlA- RFP made by CAI variants are not sufficient to explain the differences in red fluorescence in cells between different CAI (Fig. 4c, d, g, and fig. S14), and support the beneficial role of rare codons for ddlA proper fold and express, despite rare codons in transcripts being associated with mRNA destabilization and decay (38).

### Prediction of secondary structure changes in mRNA of synonymous mutants

Next, we wondered how much effect on the mRNA levels single synonymous mutants assayed in this study can have. A full exploration of synonymous mutants with RT-qPCR is not feasible due to the limited sensitivity in exploring single codon changes and the experimental cost. An alternative method to characterize the effect on the stability of mRNA transcripts is to predict their secondary structure. We predicted the secondary structure of ddlA transcripts with CoFold (39), which predicts canonical base pairs. The secondary structure of the mRNA for all possible single synonymous mutants from position 2 to 66 in the ddlA CDS (SI fig. S18) were predicted using the expected nascent transcript of ddlA mRNA (225 bases). This provides a prediction of base-pairing in the ddlA transcripts (Fig. 5a, bottom, b and SI fig. S19). The results indicate that the regions with large changes in fold efficiency upon mutation (residues 10 – 11, 20 – 22, and 51 – 54) are not associated with major disruptions in highly frequent (>80%) contacts in mRNA secondary structure (Fig. 5a, b), suggesting that secondary structure changes play a minor role in the results. Furthermore, there is no correlation between the predicted minimum free energy of mRNA transcripts and ddlA-RFP abundance (Fig. 5c, left). Additionally, RFP/GFP ratios did not reveal a difference in mRNA stability for mutants, regardless of their impact on folding efficiency or whether the mutations involved rare or common codon exchange (Fig. 5c, right and SI fig. S20).

**Figure 5.**
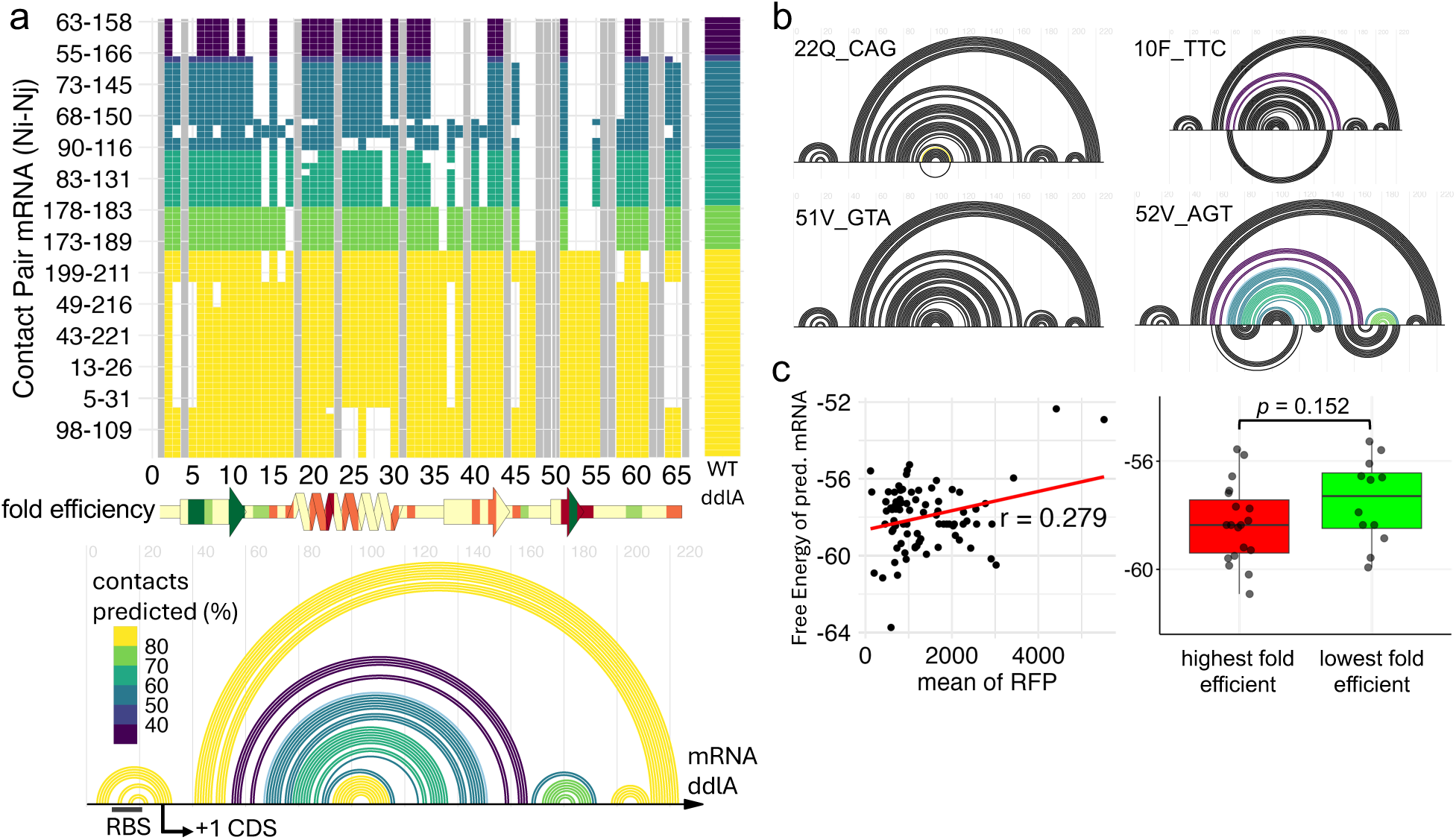
Prediction of mRNA secondary structure and minimum free energy of synonymous mutants. **a)** Heat map of the base pair contacts frequency predicted in our single synonymous mutants (fig. S18 to all possible mutants). Aligned below are the significant changes of RFP/GFP ratio depicted in the secondary structure of the ddlA protein equal to Fig 2a. At the bottom is show the base pair contacts of the WT ddlA transcript from base 1 to 225. **b)** Comparison of WT ddlA mRNA secondary structure with the mutated codon, arches in black are from the mutant prediction when exactly matches the WT in the top of the horizon and in the bottom when it does not match. **c)** Left: correlation of the predicted minimum free energy of the transcript and the measured RFP. Right: comparison of the minimum free energy of the most altered RFP/GFP ratio mutants. Comparison made with two tail t-test.

## Discussion

It is well-established that codons play a central role in regulating protein expression, folding, and function. Nonetheless, there is limited data on how choices of individual codons influence cotranslational folding. Prior work on varying codons in the CDS has involved assays for cell growth, which can be complex to interpret, or relied on specific properties of the studied gene-of- interest, such as intrinsic fluorescence (24–26, 40). Here, we present a targeted experimental approach designed to bridge this gap in a system that can be used on many different types of proteins. By targeting single-codon changes systematically in a protein domain we could investigate how folding efficiency is affected in E. coli ddlA (Fig. 1c, d).

We first hypothesized that, since ddlA is a natural E. coli gene, its native codon sequence would exhibit high folding efficiency, meaning that synonymous mutations were expected to negatively impact expression and folding. We do see a cluster of detrimental effect on the folding efficiency of synonymous mutants from positions 5 to 11 in the sequence (Fig. 2a, fig. S7), this evidence indicates that the natural codon sequence in the beginning of ddlA CDS is optimized for expression. However, we also observe that multiple mutations beyond position 15 predominantly enhance folding efficiency (Fig. 1d and Fig. 2a), with the notable exception of position 52 at the end of the β3-sheet. Overall, the emerging pattern suggests that synonymous mutations in ddlA generally improve its folding efficiency (Fig. 1d), except at the start of the CDS. A previous study demonstrated that synonymous mutations between codons in position 2 to 8 play a dominant role in modulating translation efficiency (40). The importance of this region is reflected in our mutants due to high G/C content in key codons, for instance with ^5^Arg (CGG_WT_ to AGA_mut_ and AGG_mut_), ^6^Val (GTA_WT_ to GTC_mut_ and GTG_mut_) and ^7^Gly (GGA_WT_ to GGG_mut_) which are probably hindering translation efficiency due to more secondary structures at the 5’ end of the mRNA (26, 40). Another study proposed that a “slow ramp” of rare codons at the start of the CDS enhances protein expression (41). However, in ddlA, the WT sequence consists of the most common codons in positions 2 to 13 (Fig. 4a), hence, a slow ramp during the initial translation of the CDS does not appear to be a major factor for this gene.

Mapping the folding efficiency of synonymous mutants in the secondary structure provided insights into the cotranslational folding of these elements. The α-helix appears to be more sensitive to synonymous mutations during the early stages of its decoding, as observed in three residues (^20^Ser, ^21^Leu, and ^22^Gln) in the first α-helix and with residue ^134^Ala in the α-helix connecting the C-terminal domain (Fig. 2a and 3c). The importance of codon choice for proper α-helix folding was previously observed in a study where three synonymous codon changes in the first half of a helix-turn-helix motif caused a heterologously expressed protein in E. coli to aggregate compared to the WT sequence (21). For turns connecting structured regions, we observed that most synonymous mutations affecting folding efficiency occurred immediately after the structured regions were decoded, specifically at positions 13 – 15, 30 – 32, 45 – 47, and 54 (Fig. 2c and SI Fig S7). In contrast, turns located just before β-sheets appeared to not be sensitive to codon change. A notable observation from mapping folding efficiency on β-sheets is that the last two amino acids play a critical role in determining folding efficiency, both positively and negatively (Fig. 2c).

Highlighting the most drastic effect on folding efficiency by mutants onto the tertiary structure showed important contacts of residues across the N-terminal motif, suggesting a critical role of their codons in the folding of this motif (Fig. 2c). It is known that the folding of N-terminal domain in proteins can be highly sensitive to translation kinetics (7), to the point of folding into distinct conformations in a heterologous expressed protein in E. coli after codon harmonization (42). Another study by Jiang Y. & et al. (2022) demonstrated how the folding pathway of three different proteins is affected by distinct codon patterns within their CDS. In two proteins, the altered codon usage led to a divergent state from the native structure. Interestingly, the authors discovered that codon usage induced a lasso-like entanglement in the structure of a protein (11), ddlB, a homolog to ddlA. With identical folds, it is not surprising that both proteins are sensitive to codon sequences, in contrast to a protein like DHFR that can readily refold *in vitro* (11). Given their identical topology, it is likely that a similar misfolded near-native state that is found in ddlB can also form in ddlA. These near-native states were predicted to have reduced enzymatic activity but were not expected to induce a stronger response in the PQS. Our results demonstrate a strong influence of codon choices on the PQS. Additionally, as a non-essential protein and a client of the DnaK chaperone system, ddlA may have experienced reduced evolutionary pressure to select codons that promote efficient folding.

Taken together, our data and a previous study (11) indicate that the folding pathway of ddlA is particularly prone to misfolding, possibly due to its complex topology. The data suggest that codon choice in the first half of α-helix and at the ending of β-sheets are hotspots for coordinating cotranslational folding, possibly through a decoding kinetic mechanism.

Rare codons have been observed to be enriched in α-helices and hydrogen-bonding turns (17). This agrees with our data with higher fold efficiency upon change of common to rare codons into the N-terminal helix of ddlA (Fig. 3c, ^21^Leu CTG_WT_ to CTA_mut_, and ^25^Lys AAA_WT_ to AAG_mut_). In eukaryotic ribosomes, the constriction site near the peptidyl transferase center has been demonstrated to sense the conformations of α-helices and relay that into a conformational change in the ribosome (43). Rare codons often occur in clusters in the mRNA sequence, which is expected to result in local pauses in translation and the possibility of nascent chain folding (17). Experimental evidence also highlights the functional role of codon rarity. Codons with slow translation kinetics play a critical role in E. coli protein expression and folding, as demonstrated by a study where 3 out of 8 proteins exhibited impaired expression after “slow” codons were replaced with “fast” ones (44).

We also observed a positive impact of individual rare codons over common codons on the folding efficiency of ddlA. The results show that although changes in codon rarity are consistent (Fig. 3b), there are isolated cases that do not follow this trend. For instance, in the β1-sheet at position ^10^F and ^11^G, three synonymous mutations reduced folding efficiency compared to the WT ddlA, yet one of these mutations is from a common (Gly-GGT_WT_) to rare (Gly-GGA_mut_) codon.

Changes in codon rarity can also affect folding abruptly along the sequence. In the β3-sheet, exchanges from a common to a rare codon at positions ^51^V and ^52^S resulted in opposite effects on folding efficiency (Fig. 2a and 3c). Furthermore, we confirmed the positive influence of rare codons for ddlA folding and expression with the CAI 0.3 variant made of only rare codons. Interestingly, the data for the CAI 0.3 variant is consistent with the idea that a slow translation speed allows time for the protein to fold correctly. However, the behavior of the CAI 0.3 variant conflicts with the argument regarding translation arrest due to ribosome stalling when translation is too slow (8, 45, 46).

The CAI 1 sequence was expected to result in fast translation (7, 9) but the protein abundance was low (Fig. 4c) even though its transcript level is as high as the one observed for the CAI 0.3 sequence (Fig. 4e). We further investigated the transcript levels of GFP under expression of the CAI variants to confirm that protein levels of GFP per its transcripts during expression of CAI 1 is about 40% higher than CAI 0.3 (Fig. 4f, green stripe). Perhaps the PQS response during induction of CAI 1 can offset the misfold response for translation arrest, alleviating the expression of GFP transcript and inhibiting the synthesis of ddlA-RFP (Fig. 4g). Another factor to consider is the possibility of a direct toxic effect of the CAI 1 mRNA that impairs its expression (24). Similar effect of high and low CAI sequences has been observed before, a previous study of heterologous expression in E. coli showed reduced levels of expression after optimization for CAI (CAI ≈ 1), while a less optimal sequence (CAI ≈ 0.3) had similar expression levels to the native gene sequence (47).

The effects that we observe in this study could potentially be due to changes in mRNA structure and stability, rather than cotranslational folding. The structure prediction results demonstrates that many single synonymous substitutions introduced into the sequence are likely to introduce changes in secondary and tertiary structure of mRNA (fig. S18). However, these do not correlate with the mutants that have the most substantial effect on folding efficiency (Fig. 5). Furthermore, the small sequence changes also result in very limited changes in the stability of mRNA transcripts, removing this as a major factor in explaining the result in folding efficiency. We did not quantify mRNA regulatory effects, such as the propensity of certain synonymous substitution in ddlA CDS to become targets for endoribonuclease, but these require high specificity to act so that it is unlikely to impact more than a few of the mutants in this study.

The overall methodology is an indirect assay for folding efficiency that relies on the upregulation of the stress response system in E. Coli upon production of misfolded states of proteins. While the stress reporter is associated with the DnaK promotor, DnaK itself is regulated by σ^32^. Hence, the stress signal can increase even in the absence of DnaK binding. Nonetheless, the magnitude of the response is expected to substantially increase if DnaK can bind to the misfolded protein (23). The N-terminal domain of ddlA has two DnaK binding protein sequences (fig. S8) (48), ensuring a strong response in the stress signal.

By systematically scanning synonymous substitutions in the N-terminal domain of ddlA we identified regions with both positive and negative effects on folding efficiency, highlighting where cotranslational folding is likely influenced by codon decoding kinetics. Furthermore, our findings demonstrate the role of rare codons in controlling the folding pathway in the ribosome and avoiding misfolding and how codons for residues involved in contacts in the three-dimensional space can have correlated impacts. The presented method can potentially be used to scan the CDS of recombinant proteins to provide an improved understanding of the complex process of cotranslation folding on the ribosome and the evolution of codon sequences. Our results also highlight the complex relationships between codon rarity and sequence position, secondary structure and topology in cotranslational folding, and the challenges in establishing simple models to explain these phenomena.

## Materials and Methods

### Cloning of AraBAD Promoter in the Reporter Plasmid

The plasmid containing the araBAD promoter, pBAD-sfGFP (Addgene Plasmid #85482), was used as the source of the promoter sequence. The araBAD promoter region, including araC, was PCR- amplified using a forward primer with a SpeI restriction site at the 5’ end and a reverse primer with a EcoRI site in the 5’ end (SI Table S1). The nucleotide coding sequence of ddlA (NCBI Reference Sequence: NP_414915.1) from E. coli K-12 addressed as wild-type, was synthesized by GENEWIZ® from Azenta and ordered cloned inside the original plasmid used as stability sensor (containing the T7 promoter). To remove the original promoter, the plasmid needed to be mutated at bases -10 and -12 upstream the promoter region to introduce a SpeI cleavage site following the protocol described in the next section. Mutagenesis was confirmed by incubating the miniprepped plasmid (QIAprep Spin, QIAGEN) with SpeI (FastDigest, Thermo Scientific™), followed by agarose gel electrophoresis using GelStar® (Lonza Bioscience) staining to visualize linearized plasmid bands. After verification, the amplified araBAD promoter and the mutated reporter plasmid were digested with SpeI and EcoRI (FastDigest, Thermo Scientific™), gel-purified, and ligated using T4 DNA ligase (Thermo Scientific™), following the manufacturer’s recommendations. To remove LacI and additional chloramphenicol resistance from the reporter plasmid, it was digested with SwaI and FspAI, generating a blunt-ended DNA fragment. The digestion product was gel- purified, and the main plasmid fragment was ligated using T4 DNA ligase. The final plasmid constructs were transformed into competent E. coli Top10 cells for amplification miniprepped. Whole plasmid sequencing (ONT lite sequencing, Eurofins Genomics) was performed to confirm the correct assembly of the final construct and the sequence of the ddlA WT. Synonymous mutants were made with miniprepped purifications of the final construct.

### Single Synonymous Mutagenesis Scan

The nucleotide coding sequence of ddlA WT was used as input in a modified python script (49) to design primers containing all the possible synonymous counterpart for every position. Then, pairs of mutated primers are sorted by melting temperature of 72 °C to select the most stable. For positions containing amino acids that only have one synonymous codon substitution (*e.g*., Asn, Asp, Cys, Gln, Glu, His, Lys, Phe and Tyr), a single complementary pair of forward and reverse primer containing the substitution was selected (SI Table S3). For the remaining amino acids with more than one possible codon substitution (*e.g*., Ala, Arg, Gly, Ile, Leu, Pro, Ser, Thr and Val) a forward primer for one codon substitution was paired with the corresponding reverse for other substitution (SI Table S3). The reason to not have complementary pairs for all codons was to reduce the cost associated with acquisition of primers and number of reactions. Mutagenesis was made following stablished site-direct mutagenesis (50). Each reaction consisted in three steps, first plasmid elongation using separated primers, second, pooling complimentary reactions together and submitting for stepwise cooling from 95 °C to 37 °C, and third, adding DpnI to degrade template plasmid. Then the reaction mixture is transformed into E. coli strain BL21 for selection in LB-agar with Ampicillin (50 µg/ml) and Chloramphenicol (25 µg/ml). At least four isolated clones are single picked with sterile wood toothpicks and inoculated in 300 µl of TB with both antibiotics in a 96- deep well plate for 15h at 35 °C. Finally, cell cultures are diluted 5 times in LB and 5 µl are transferred to a commercial pre-paid LB-agar PlateSeq kit Clones–Ampicillin (Eurofins Genomics, Product n° 30PK-000PPC) for sequencing. Another 5 µl from the same diluted culture are transferred to a 96-microtiter plate with LB-agar with antibiotics as record of cells for later analysis of confirmed synonymous mutants. Both plates are incubated for 15h at 37 °C. Then, the PlateSeq kit is sent for sequencing using rolling circle amplification to confirm the presence of the expected single synonymous mutants with a primer designed to anneal 100 bp upstream of the second codon of ddlA (SI Table S1).

### Codon Rarity and ddlA Single Frequency Codons Variants

Codon sequence for the unique rare and common were generated based on the lowest and highest codon frequency per amino acid based on the codon usage table from Kazuka database (36) for Escherichia coli strain K12 (taxon ID: 83333). Codon rarity was based on Kazuka table too, rare codon category was made of codons with frequency less than 11 per thousand, which include 23 codons, while common codons are the remaining 38 possible codons. CAI values were calculated using web version of CAIcal (51). Full sequences were ordered already in the report plasmid through onboard cloning option with GENEWIZ® from Azenta.

### Expression of ddlA-RFP and flow cytometer analysis

Clones containing the unique and positive synonymous mutant are identify by a python script from the clipped sequences provided by Eurofins Genomics. From the plate record, each confirmed synonymous mutant are picked and inoculated in three different wells, to account for biological triplicates, in a 96-deep well plate with 300 µl of TB (with antibiotics) for 15h at 35 °C. To synchronize cell growth for the inoculum, the plate is divided in 4 time slots of initial inoculum with a time difference of 50 minutes. Then, cells are diluted 5 times in a 96-PCR plate and 10 µl of each is transferred to a 700 µl of thermalized LB media pH 7 at 37 °C with antibiotics for grow in an orbital shaker at 1100 rpm until all the wells have reached OD_600_ = 0.350 measured in a 96- microtiter plate containing 200 µl of culture. This is followed by induction of expression by mixing 1:1 (V:V) the inoculum with LB containing 0.2% of L-arabinose thermalized at 37 °C to yield a final concentration of 0.1%. Then, cell culture is incubated at 37 °C in an Inheco shaker with 2 mm side to side amplitude at 13.5 times/second. After ∼3h of induction and measured OD_600_ > 0.400 in a 96-microtiter plate containing 200 µl of culture, 15 µl of each cultured well is transferred to 1 ml of PBS at room temperature and analyzed by the S3e cell sorter from BIO-RAD with a flow rate of approximately 2000 events/seconds. Red and green fluorescence, as well as RFP/GFP ratios readouts, correspond to the median intensity of 10,000 E. coli events. Fluorescence was detected using a 525/30 filter (green fluorescence) and a 615/25 filter (red fluorescence). The mean and standard deviation from three biological replicates were used for data analysis. Flow cytometry data were analysed using the flowCore and flowViz packages within the Bioconductor framework (52). The growth rate was calculated as the difference in the natural logarithm of OD₆₀₀ measured from cells just before dilution in PBS for flow cytometry analysis and at the moment of induction, divided by the elapsed time between them.

### Real Time – qPCR Assay

Cells were grown in a 96-deep well plate and induced as described in the previous section. mRNA extraction was done with PureLink™ RNA Mini Kit from Invitrogen™, according to the manual, after 4h of induction of L-arabinose. About 0.6 µg/µl of purified mRNA of each sample was treated with 3 Units of DNase I Amplification Grade from Invitrogen™ following manufacture’s recommendations. All RT-qPCR reaction were made in a final volume of 15 µl using Luna® Universal One-Step RT-qPCR Kit from New England Biolabs® in a thermocycle CFX96™ Real-Time System from BIO-RAD with the following program: 55 °C for 10 minutes, 95 °C for 1 minute, then 40 cycles of 10 seconds at 95 °C and 25 seconds at 60 °C recording the fluorescence, then melting curves from finished reactions were made from 60 °C to 90 °C increasing the temperature by 0.5 °C/cycle and each cycle during 5 seconds. Relative gene expression was calculated using 2−ΔΔCT values and validated (53) for two reference genes, cysG and idnT (54) (fig. S16). PCR amplification curves from experiment in Fig. 4d–f is shown in fig. S15. Confirmation of amplified amplicons in purified mRNA is shown in 1.2% agarose gel in fig. S17. Primers used for RT-qPCR experiment are listed in SI Table S2.

### Secondary structure prediction of mRNA

Based on the AraBad promoter region –35 and –10 proximal to the ddlA WT CDS, the transcript starting from the next nucleotide +1g until the codon referent to position 66 was used as input to predict its secondary structure using the software CoFold (39). mRNA prediction of single synonymous mutants in the CDS region were made for every possible mutant. Arc diagrams of RNA secondary structure were made with R-chie (55) using the predicted mRNA structures.

### Statistical analysis

Difference between the mean of two groups of samples were evaluated with two tailed t-test if their data were confirmed to be normally distributed by Shapiro-Francia test. Data without normal distribution used Wilcoxon rank-sum test to evaluate significant differences between the median of two defined groups.

## Acknowledgments

We are grateful to Camille Wernersson for insightful discussions and valuable ideas regarding cloning and site-directed mutagenesis strategies. We also appreciate the assistance of Hélène Bret in providing scripts for automating the selection of primers for mutagenesis experiments. Additionally, we thank Dominique P. Fastus for his helpful discussions on mRNA secondary structure prediction.

## Author contributions

Writing—original draft: F.A.M.O.

Writing—review & editing: F.A.M.O., I.A.

Conceptualization: I.A.

Resources: I.A.

Methodology: F.A.M.O., I.A.

Data curation: F.A.M.O.

Investigation: F.A.M.O., I.A.

Visualization: F.A.M.O.

Funding Acquisition: F.A.M.O., I.A.

Supervision: I.A.

## Data and materials availability

All data are available in the main text or the supplementary materials.

