## Supplementary Material for "Effects of single synonymous substitutions on folding efficiency demonstrate the influence of rare codons and protein structure"

Felipe Akihiro Melo Otsuka & Ingemar André*

**This PDF file includes:**

Figs. S1 to S20

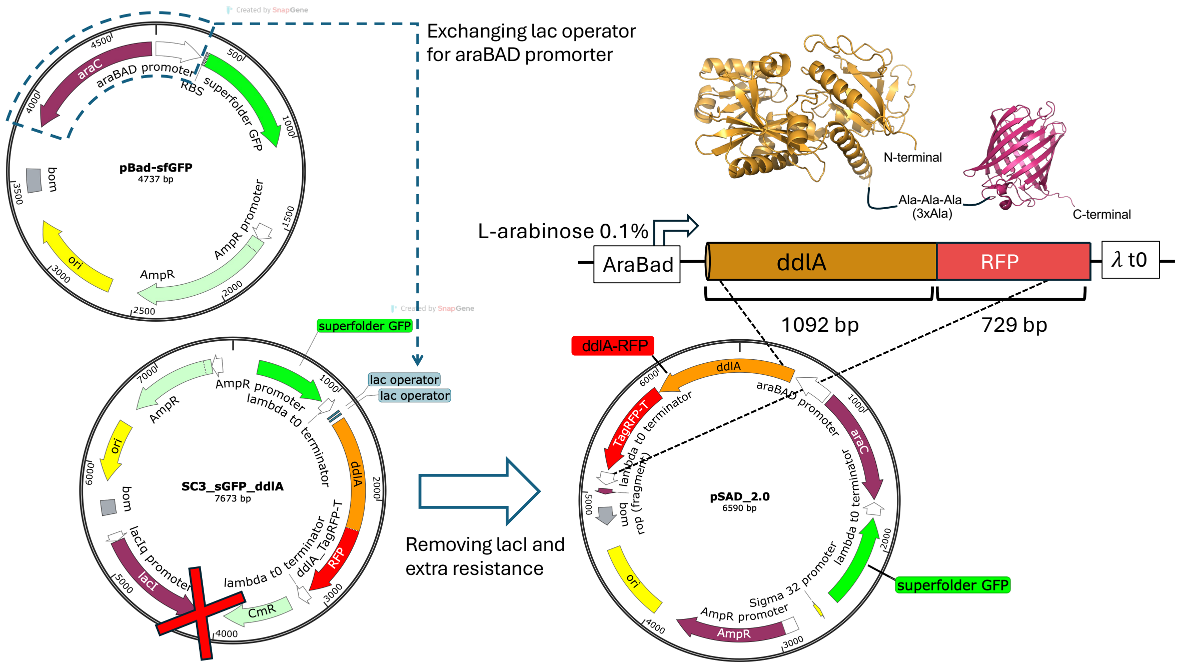


**Fig. S1.** The AraBAD promoter from pBAD-sfGFP (Addgene #85482) was PCR amplified using primers (Table S1) and replaced the T7 promoter from the previous SC3_sfGFP_ddlA reporter plasmid to construct pSAD_ddlA-RFP_sfGFP. The LacI promoter and chloramphenicol resistance gene were removed by digestion with two blunt-end restriction endonucleases, SwaI and FspAI, while ampicillin resistance was retained.

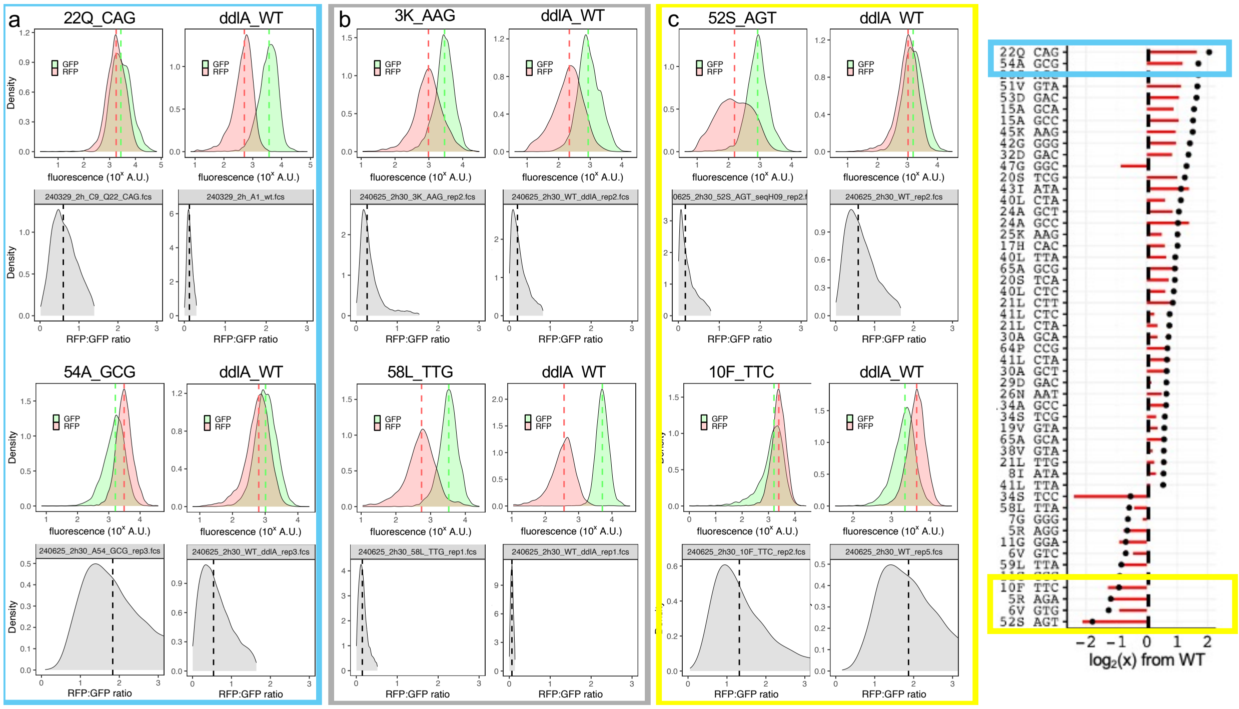


**Fig. S2.** Histograms of six single synonymous mutants, each paired with WT ddlA, which was used for median normalization. a) Two mutants highlighted in the blue box in the rightmost graph showed the greatest improvements in folding efficiency. b) Two mutants in the grey box were neutral regarding folding efficiency. c) Two mutants in the yellow box were the most detrimental to folding efficiency.

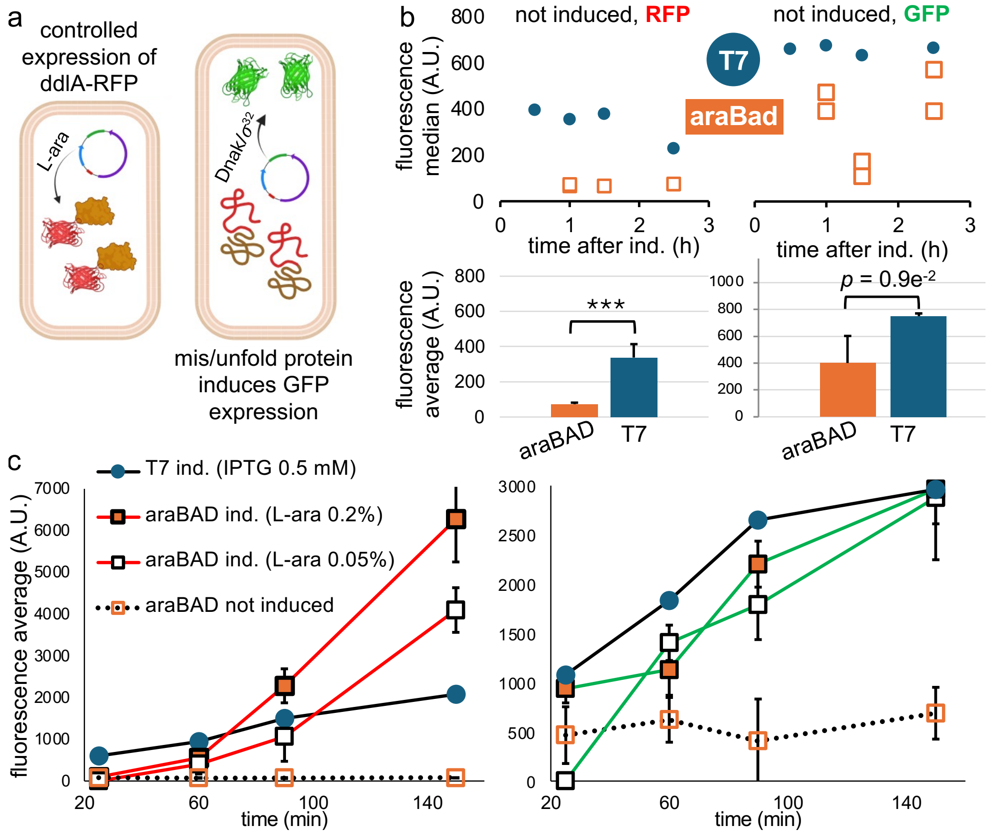


**Fig. S3.** Controlled expression of ddlA-RFP with AraBAD promoter and comparison with the T7 promoter. a) Schematic representation of ddlA-RFP expression induced by L-arabinose, with GFP serving as a reporter for DnaK/σ32 activation in E. coli. b) Fluorescence measurements of non-induced E. coli cells analyzed by FACS. The left panel shows red fluorescence (RFP), while the right panel shows green fluorescence (GFP), comparing plasmids with either T7 (blue circles) or araBAD (orange squares) promoters. c) Expression in Erlenmeyer flasks (20 mL of liquid cultures) with fluorescence measurements taken over 150 minutes post-induction. Induction was performed at 37 °C, and values represent the averages of biological triplicates. The araBAD promoter provides a key advantage in tightly regulating ddlA-RFP expression, effectively minimizing background expression prior to induction (b). At 60 minutes post-induction, the total accumulated ddlA-RFP and GFP levels remain higher under the T7 promoter (c, left graph). However, by 90 minutes, cells induced with 0.2% L-arabinose (araBAD promoter) surpass T7-driven expression in ddlA-RFP levels (c, left graph), while accumulated GFP fluorescence remains lower (c, right graph). This suggest that expression leakage is being accounted for the GFP expression until 90 minutes of induction. Only after 150 minutes do both promoters result in similar GFP levels (c, right graph), yet ddlA-RFP production under the araBAD promoter remains at least twofold higher (c, left graph). This suggests that more properly folded protein is produced per unit of PQC activation (GFP signal) reinforcing the role of a controlled transcription from araBAD promoter to prevent higher GFP background.

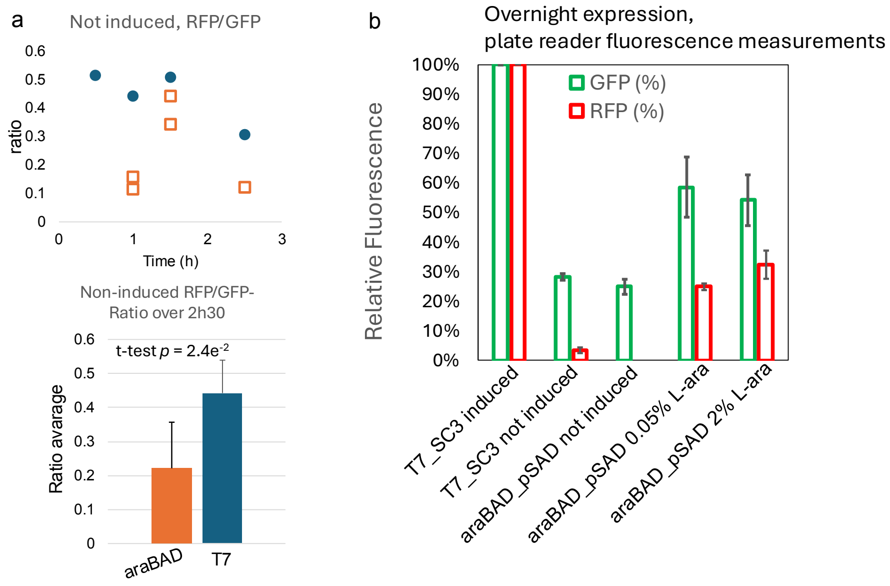


**Fig. S4**. a) Ratio of RFP/GFP in non-induced E. coli carrying pSAD_ddlA-RFP_GFP plasmid. b) Relative overnight expression of ddlA-RFP, comparing the older plasmid with a T7 promoter (induced by IPTG) and the newer plasmid with an araBAD promoter (induced by L-arabinose).

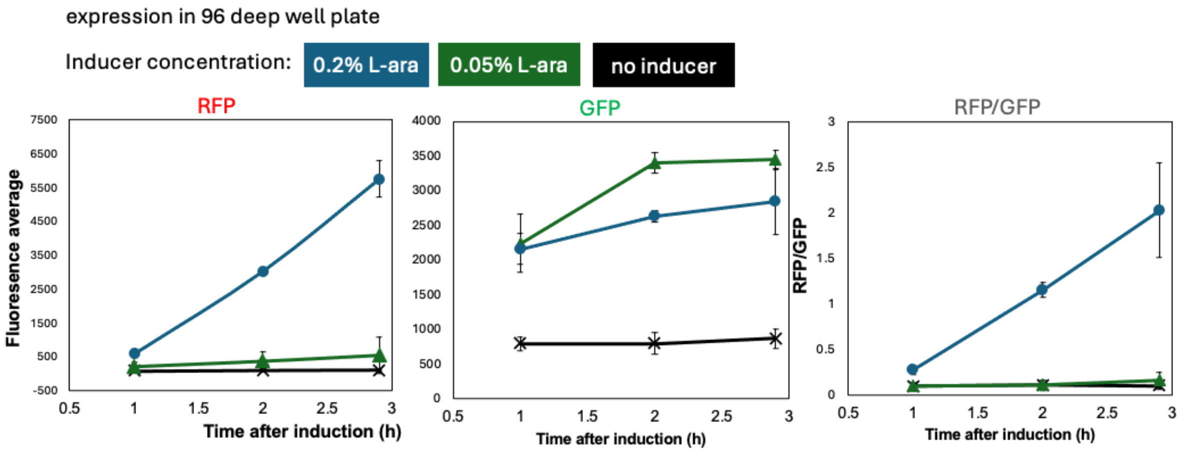


**Fig. S5**. a) Fluorescence measurements of cells carrying pSAD_ddlA-RFP_GFP with different concentration of L-arabinose as inducer. RFP and GFP readouts in three times post-induction, at 1h, 2h and 2h55. Expression of cells made in 700 µL of liquid culture in 96-deep well plate. Values are means of fluorescence median from three replicates.

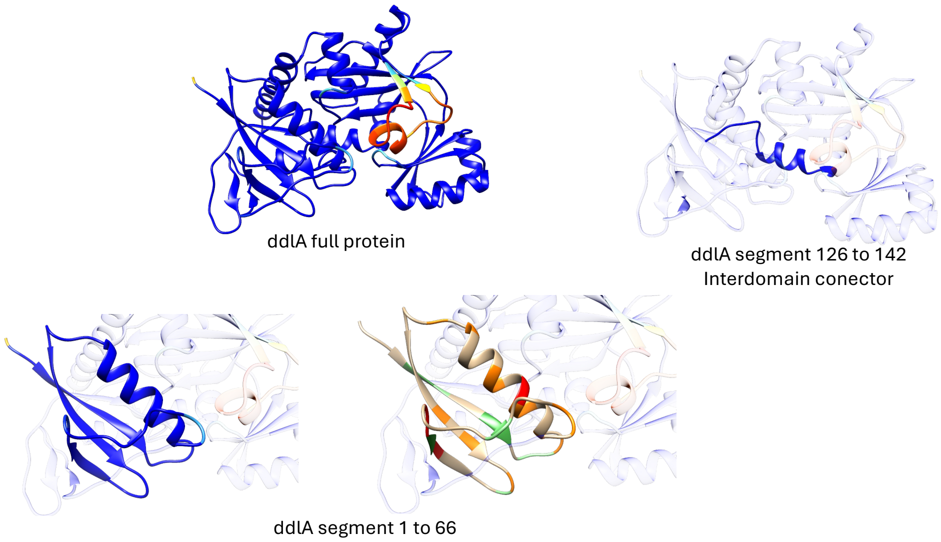


**Fig. S6**. AlphaFold2 prediction of ddlA colored by standard pLDDT. Highlighted in solid colors on the structural segments where synonymous mutants where made is shown.

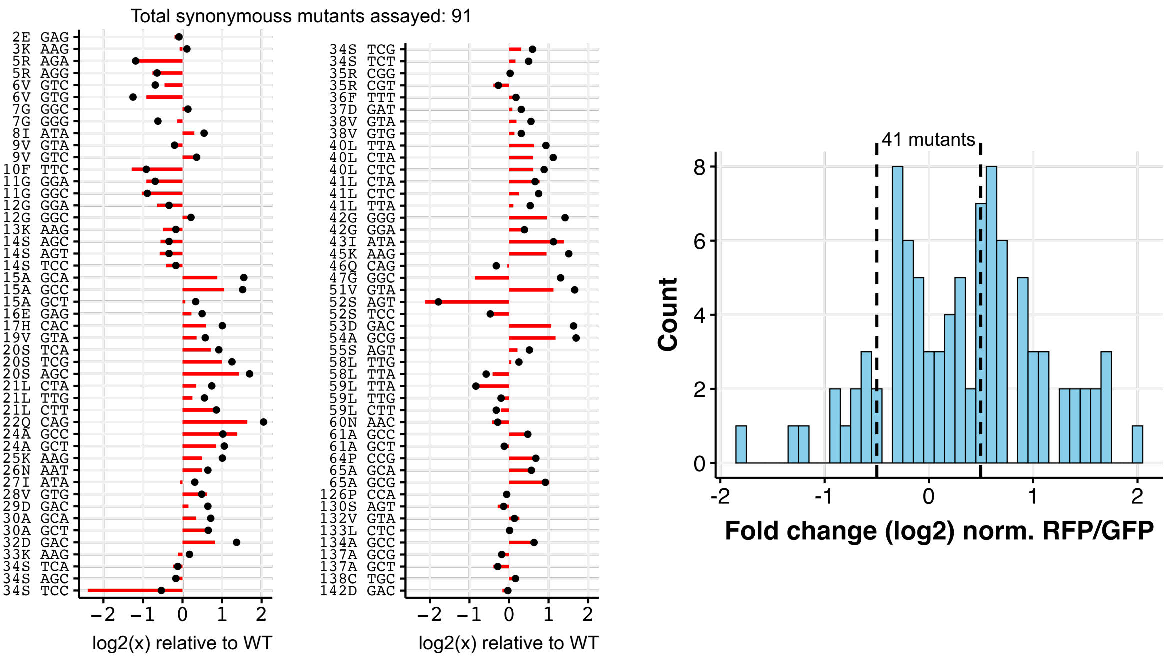


**Fig. S7**. FC ratios of each mutant (black dots) and RFP levels (red bars) compared to the ddlA WT for all confirmed and analyzed single synonymous mutants in this study. the histogram on the right displays the distribution of normalized FC ratios for all the mutants.

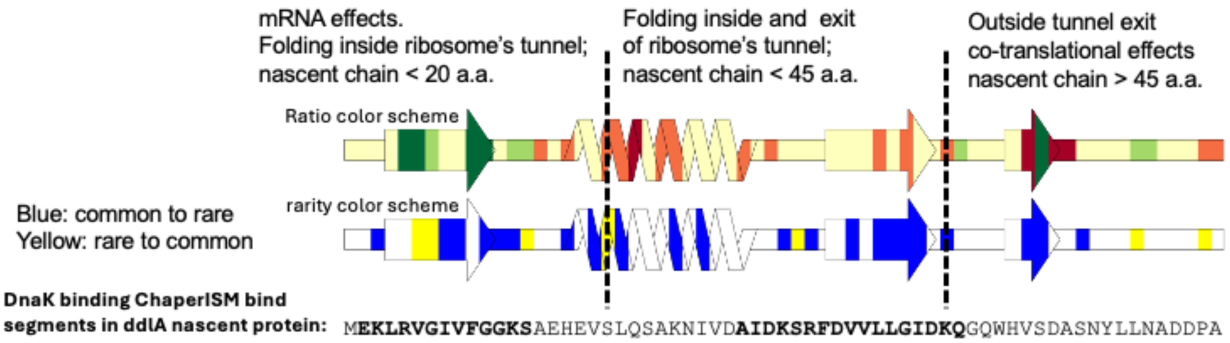


**Fig. S8**. Schematic representation of ddlA's secondary structure with annotated RFP/GFP measurements, as shown in Fig. 2, along with changes in codon rarity. The diagram also indicates the expected thresholds where the nascent polypeptide chain is positioned during ribosomal translation. At the bottom, the aligned protein sequence highlights in bold the predicted Dnak binding segments.

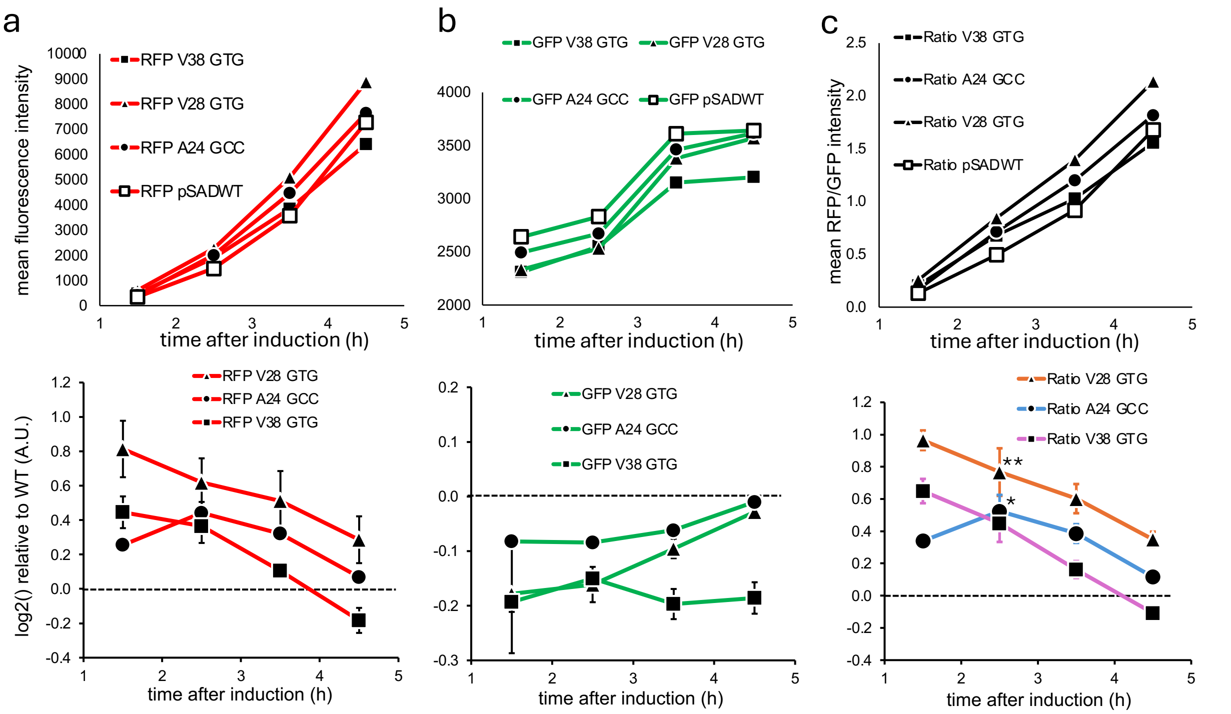


**Fig. S9**. Kinetics of ddlA-RFP expression from 1 to 4.5 hours half after induction with L-arabinose in Erlenmeyer flasks (20 mL culture). Wild type ddlA (pSADWT, open squares) and its synonymous mutants $^{24}$Ala (GCC) (circles), $^{28}$Val (GTG) (triangles and $^{38}$Val (GTG) (filled squares) were expressed in three biological replicates. **a**, **b**) Mean fluorescence of RFP and GFP, followed by normalization with the WT values. **c**) Mean RFP/GFP ratios and normalized ratios of synonymous mutants. Mutant $^{28}$Val (GTG) (t-test, $p < 0.01$) and $^{24}$Ala (GCC) (t-test, $p < 0.05$) showed significant differences from the WT after 2.5 hours of induction.

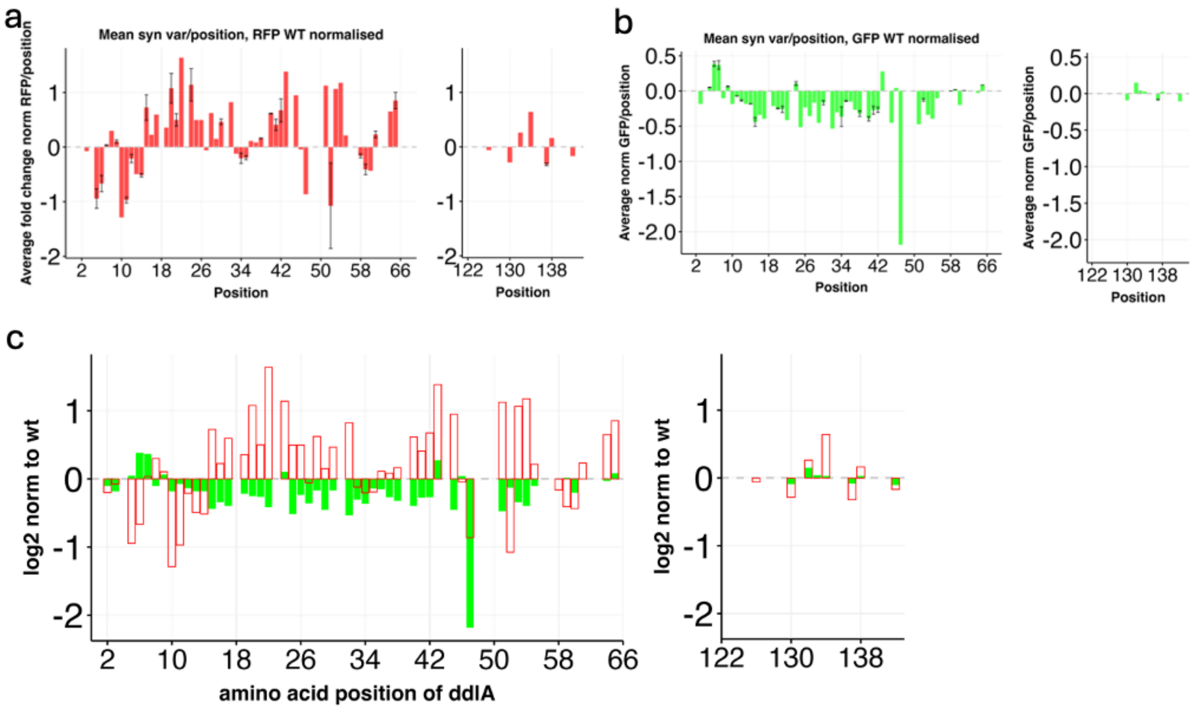


**Fig. S10**. **a**–**c**) Per-position average of fold change of RFP and GFP readouts for synonymous mutants relative to the WT ddlA CDS. Standard deviations are calculated only for positions where more than one mutant was assayed.

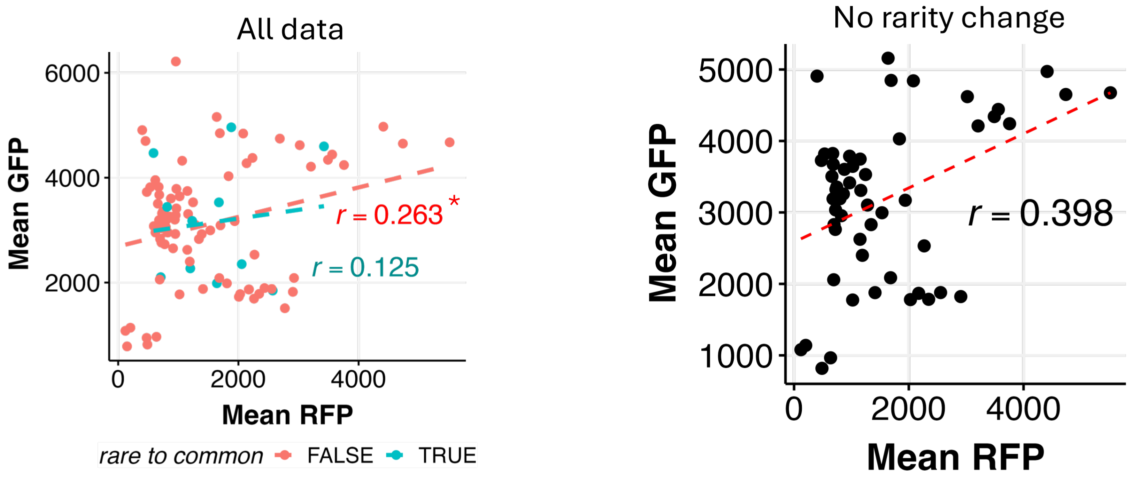


**Fig. S11**. Pearson correlation of mean fluorescence measurements. The left panel shows data categorized by changes in codon rarity, while the right panel isolates cases where no change in codon rarity was made.

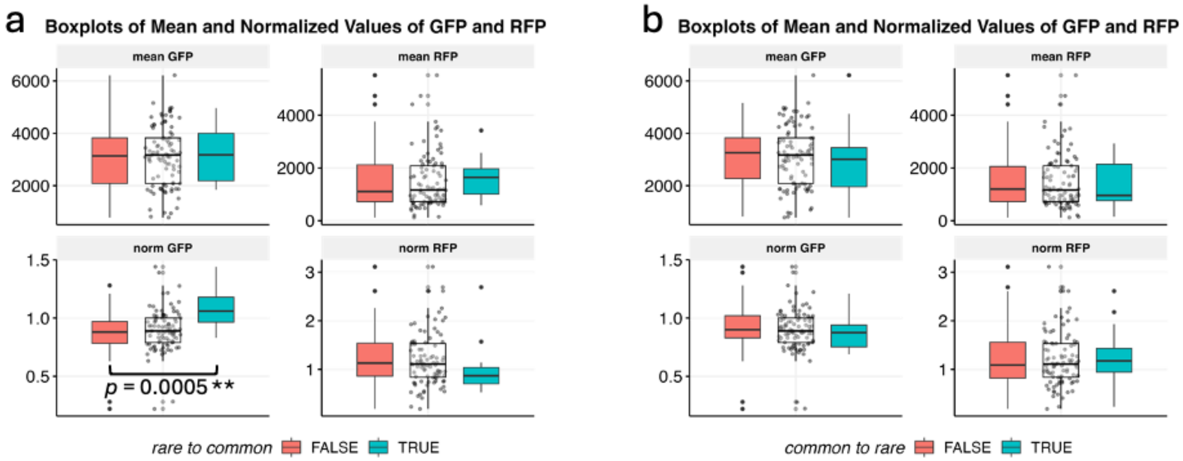


**Fig. S12**. Box plots of RFP and GFP values categorized by changes in codons from rare to common (**a**) and common to rare (**b**): *FALSE* (red) and *TRUE* (blue). Non-categorized data is represented by the white box in the middle. The top panels show the mean values, while the bottom panels display values normalized to the WT. Only the significant difference ($p < 0.05$) is shown after testing with Wilcoxon rank-sum test between *TRUE* and *FALSE* categories.

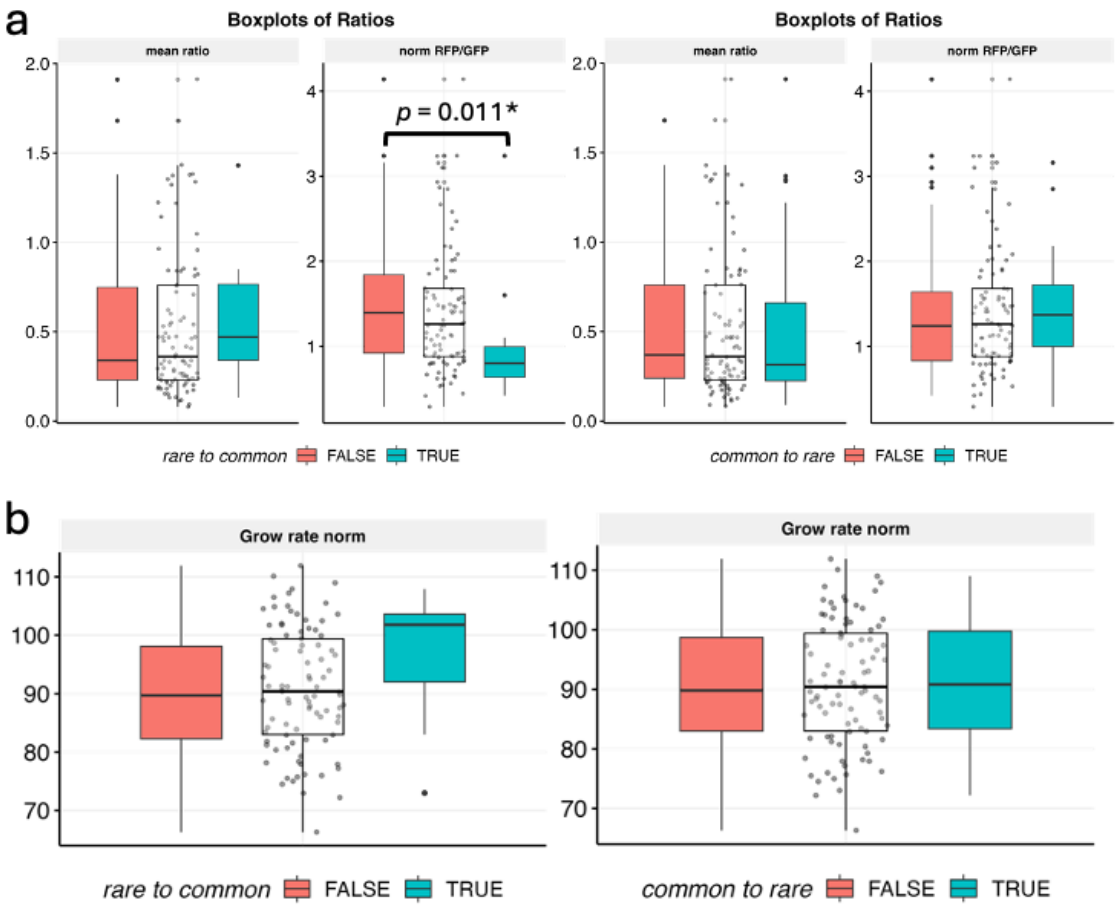


**Fig. S13**. **a**) Box plots of RFP/GFP mean and normalized (by the WT) values categorized by changes in codons from rare to common (left) and common to rare (right): *FALSE* (red) and *TRUE* (blue). **b**) Box plots of normalized growth rate compared to the WT. Non-categorized data is represented by the white box in the middle. Only the significant difference ($p < 0.05$) is shown after testing with Wilcoxon rank-sum test between *TRUE* and *FALSE* categories.

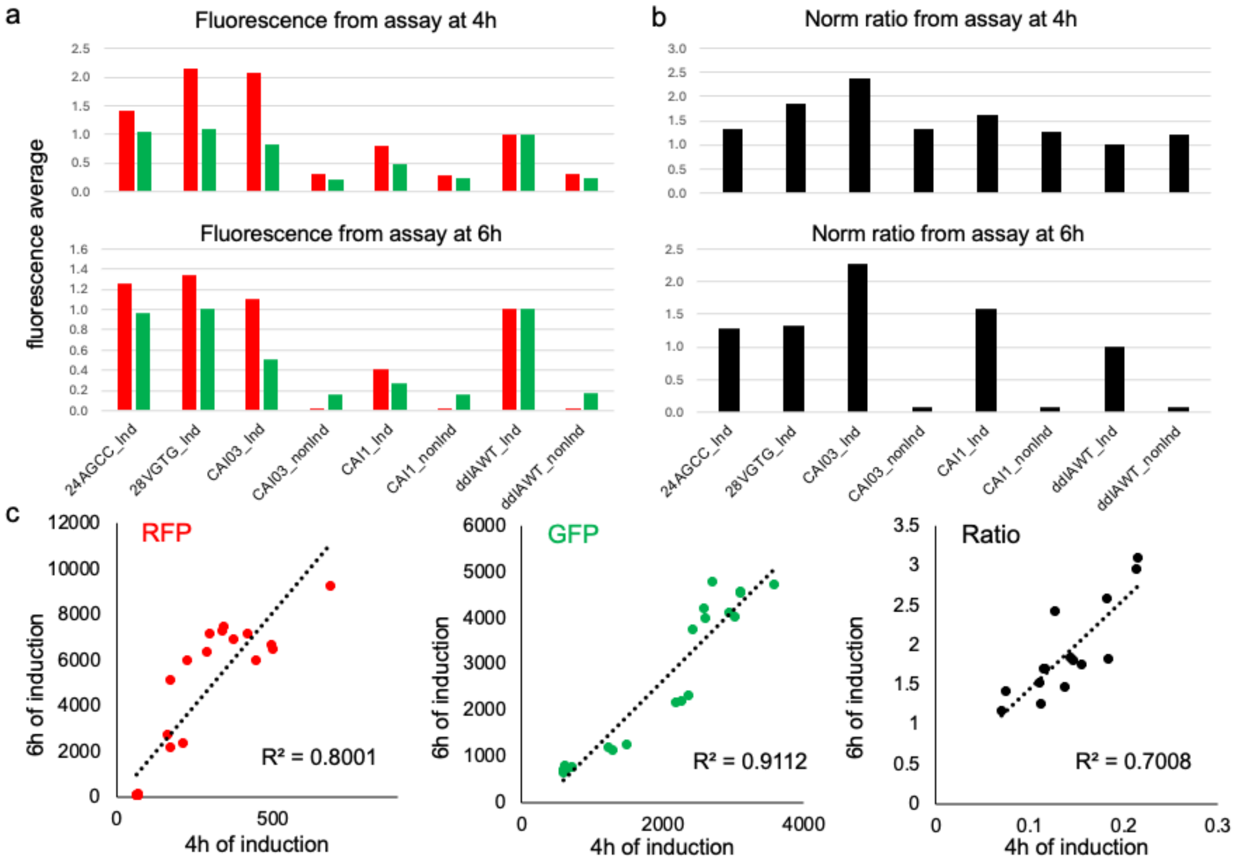


**Fig. S14**. Flow cytometry fluorescence measurements of induced and non-induced ddlA-RFP cells in a 96-deep well plate intended for RT-qPCR analysis. **a**) Normalized fluorescence by the WT ddlA, RFP (red bars) and GFP (green bars) readouts of cells at 4 hours (top) and 6 hours (bottom) post-induction with L-arabionse. **b**) Normalized RFP/GFP ratios relative to WT ddlA after 4h (top) and 6h (bottom) after induction. Data represent the mean of biological triplicates. **c)** Correlation of fluorescence readouts (RFP on left, GFP on middle) and ratio (right) at 4 hours and 6 hours post-induction of WT ddlA, $^{24}$Ala (GCC), $^{48}$Val (GTG), CAI 0.3 and CAI 1. The observed linearity suggests uniform expression and a consistent stress response even after prolonged induction in single synonymous mutants and CAI variants. Data represent mean of biological triplicates.

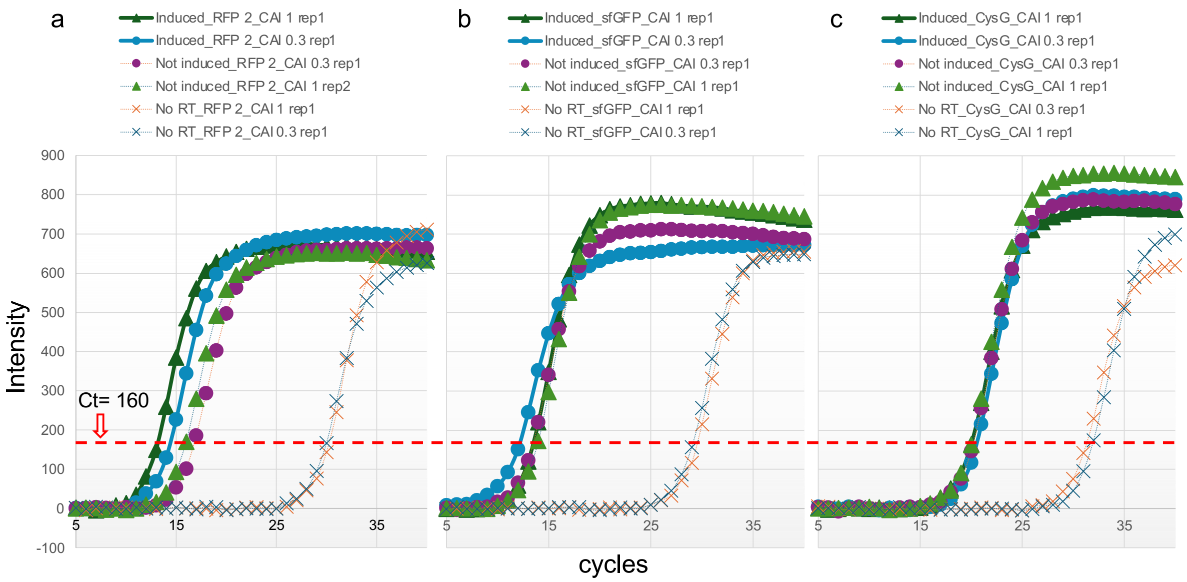


**Fig. S15**. RT-qPCR amplification curves targeting ddlA-RFP transcripts (RFP2 primers) in “a”, sfGFP in “b” and CysG (reference gene) in “c”, corresponding to the data present in Fig. **3d–f**. Each plot shows a single biological and technical replicate of samples. Samples include cells induced and non-induced controls at 4 hours post-induction of CAI 0.3 CAI 1 variants and WT ddlA. Reaction controls without reverse transcriptase (No RT_) are also shown.

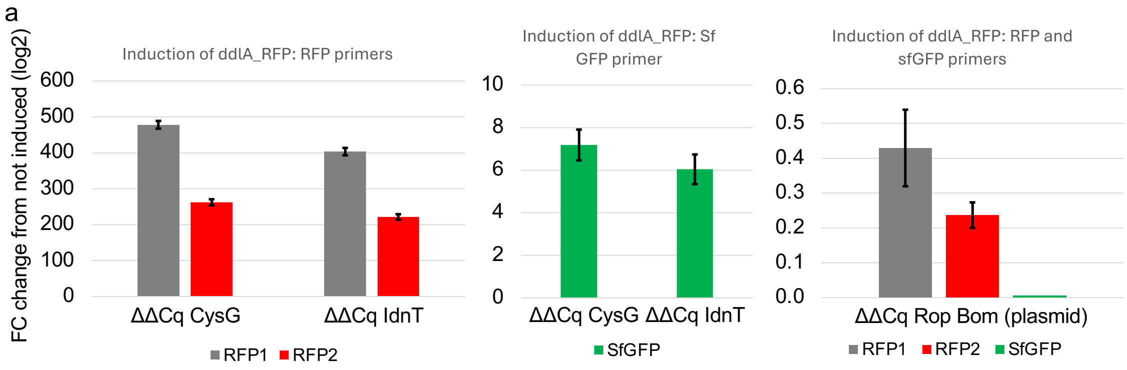


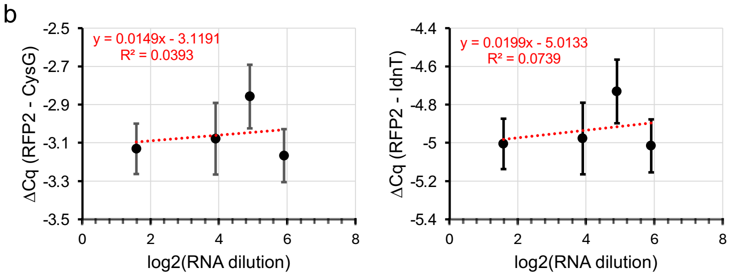


| No Reverse Transcriptase Cq Mean | No RT STDEV |
|---|---|
| 30.5 | 1.5 |

**Fig. S16**. Overall RT-qPCR validation. a) Fold change in expression levels relative to non-induced cells carrying pSAD_ddlA-RFP_sfGFP, induced with L-arabinose (0.1%). Primers pairs targeting the RFP transcript (RFP1 and RFP2, Table S2) and sfGFP transcripts. CysG and IdnT refer to primer pairs targeting reference genes in the E. coli. “Rop Bom” refers to a primer pair targeting the plasmid DNA, serving as a negative control. b) Compared efficiency of Ct values for the RFP2 primer pair and reference genes (CysG and IdnT). Four dilutions were assayed, spanning 20-fold range from the most concentrated to the most diluted sample of purified RNA. The table (right) display the mean Ct value and standard deviation (SD) for reactions performed without the reverse transcriptase (No RT control).

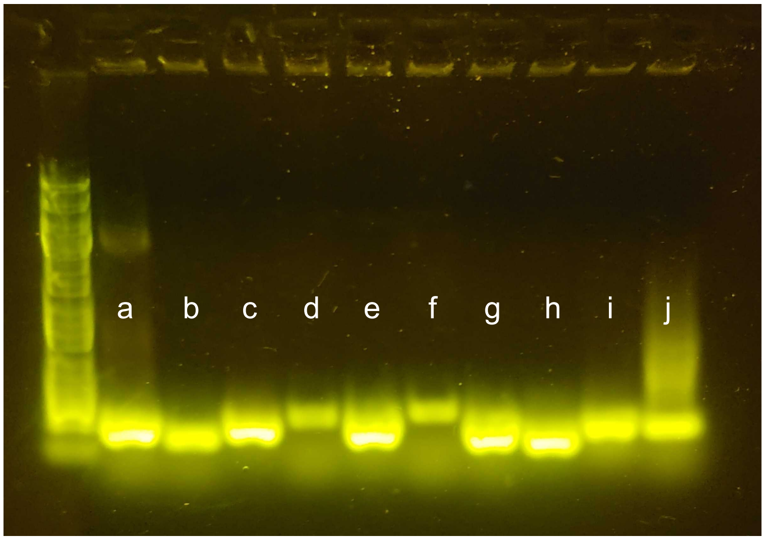


**Fig. S17**. Reverse transcriptase PCR reactions in agarose gel with plasmid as control (a, b) and RNA extracted from cells induced with L-arabinose (c – j). a) pSAD_ddlA-RFP_sfGFP with primers RopBom; b) pSAD_ddlA-RFP_sfGFP with primers RFP2; c) Induced cells with RopBom; d) Induced cells with RopBom without reverse transcriptase; e) Induced cells with primers RFP2; f) Induced cells with primers RFP2 without reverse transcriptase; g) Induced cells with primers sfGFP; h) Iinduced cells with primers rrsA; i) Induced cells with primers CysG; j) Induced cells with primers IdnT.

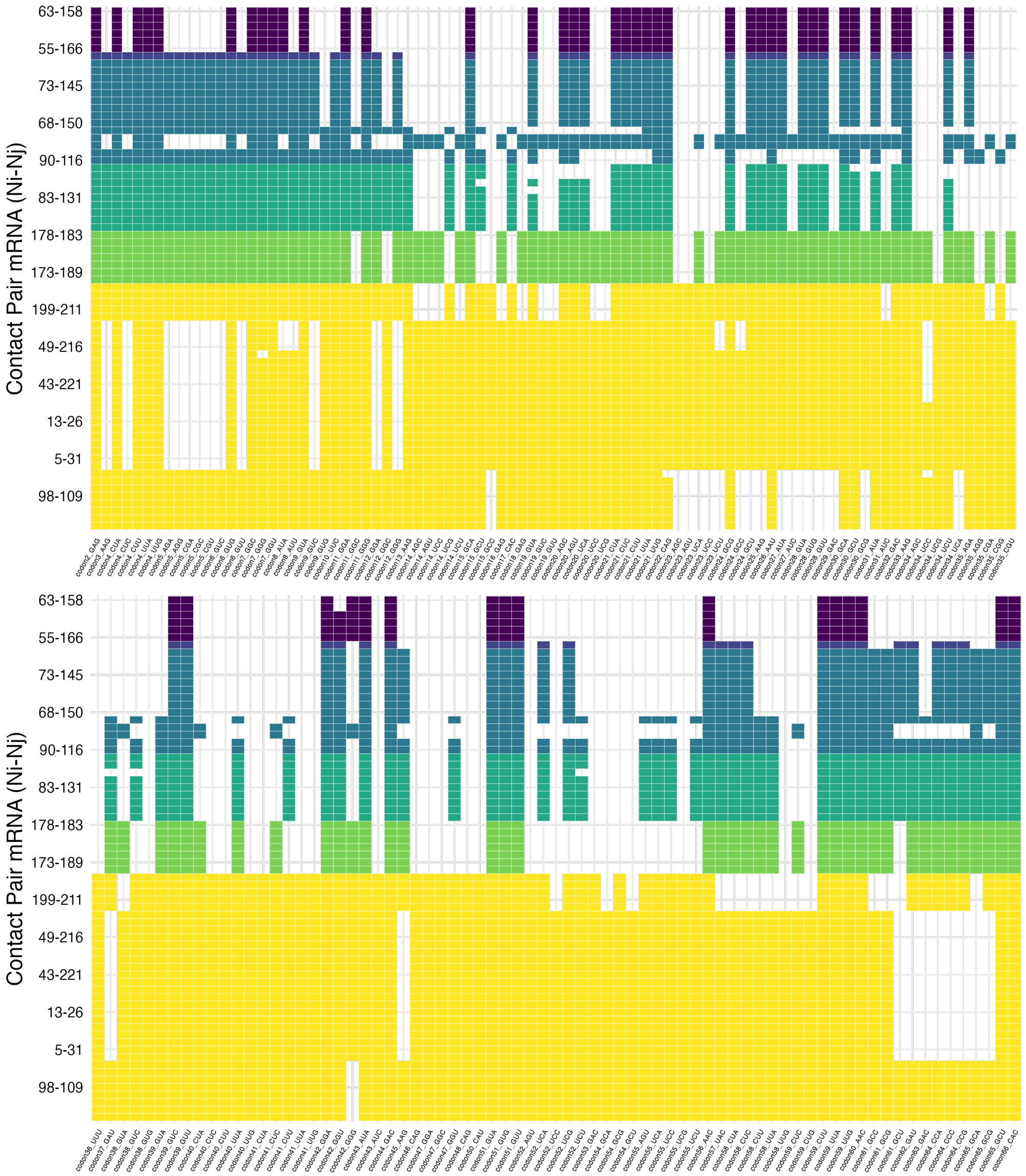

**Fig. S18**. Compiled base pair frequency of all possible single synonymous mutants of ddlA from positions 2 to 66. Base pairs with a frequency greater than 80% across all predicted secondary structures are highlighted in yellow. Detailed color scheme is displayed in **Fig. 5a** of the main text.

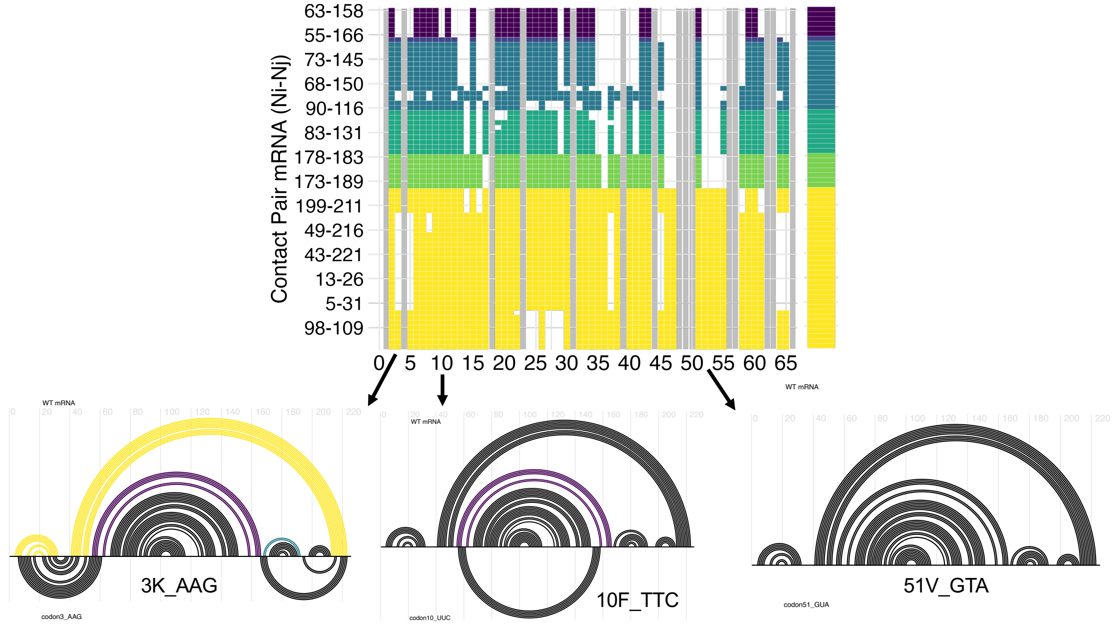


**Fig. S19**. Additional examples of mRNA secondary structure predictions for mutants. Arch plots illustrate predicted structural differences compared to the WT mRNA. Upper-yellow-colored arches indicate newly formed contacts that differ from the WT structure, while fully upper black arches represent contacts that remain unchanged between the mutant and WT transcript.

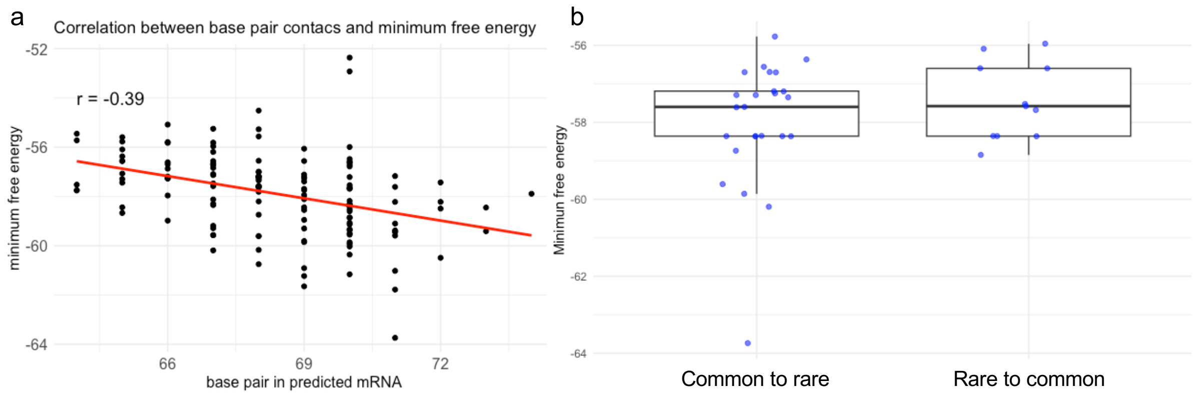


**Fig. S20**. a) Correlation analysis between the predicted minimum free energy and the number of base pair contacts in mRNA secondary structure. b) Comparison of predicted minimum free energy values between two codon exchange categories: common to rare and rare to common.

**Table S1.** List of primers used to clone the promoter AraBAD in plasmid reporter SC3_sGFP_ddlA and for sequencing of synonymous mutants.

| Primer name | Sequence - 5' to 3' | Target plasmid | Purpose |
|---|---|---|---|
| Mut_ttt1094cta | **F**w: aacaccactagtcgagaaatcataaaaaatttatttg<br>Rv: atttttatgatttctcgactagtggtgtt | SC3_sGFP_ddlA | Mutating TT1094CTA for RE SpeI (AˆCTAGT) to cut upstream T7 promoter. |
| araBAD_SpeI_Eco | **F**w: agcagACTAGTttatgacaacttgacggctacatc<br>Rv: gaatcGAATTCgtatggagaaacagtagagagttg | pBAD_sfGFP | Amplify araBAD promoter with 5'-SpeI and EcoRI-3'. |
| ddlA_seq4 | Fw**:** gtcacactttgctatgccatag | pSAD_ddlA-RFP | Anneal 101 bases upstream the second codon from ddlA for sequencing. |

**Table S2.** List of primers for RT-qPCR.

| Target | ID | Primer sequence | Amplicon size |
|---|---|---|---|
| ddlA-RFP | RFP1 | Fw: atcaaggaggccgacaaagag<br>Rv: cttgtacagctcgtccatgc | 177 |
| ddlA-RFP | RFP2 | Fw: tgcagaagaaaacactcggc<br>Rv: tgtcggcctccttgattctttc | 222 |
| sfGFP | sfGFP | Fw: gttcactggtgtcgtccctattc<br>Rv: gagcaaagcactgaacacc | 197 |
| CysG | CysG | Fw: gctggcaaacaacgatcaga<br>Rv: gtagaccaccacatctgcctg | 169 |
| IdnT | IdnT | Fw: gccagtaccattaaagcggg<br>Rv: tcgccgttattgatgtgctg | 143 |
| pSAD | RopBom | Fw: cctctgacacatgcagctcc<br>Rv: catgtggtgcactctcagtacaatctg | 201 |
| rrsA | rrsA | Fw: cgatccctagctggtctgag<br>Rv: ttcttcatacacgcggcatg | 136 |

**Table S3.** List of primers for site-directed mutagenesis. Rows corresponds to the pair of primers used in the mutagenesis reaction to obtain a synonymous mutant.

| Forward primers | Sequence - 5' to 3' | Length | Reverse primers | Sequence - 5' to 3' | Length |
|---|---|---|---|---|---|
| 2E_GAG | gaggagaaattaactatggagaaactgcgggtaggaatcg | 40 | 2E_GAG | cgattcctacccgcagtttctccatagttaatttctcctc | 40 |
| 3K_AAG | ggagaaattaactatggaaaagctgcgggtaggaatcg | 38 | 3K_AAG | cgattcctacccgcagcttttccatagttaatttctcc | 38 |
| 4L_CTT | gaaattaactatggaaaaacttcgggtaggaatcgtttttg | 41 | 4L_CTT | caaaaacgattcctacccgaagtttttccatagttaatttc | 41 |
| 4L_CTC | gaaattaactatggaaaaactccgggtaggaatcgtttttg | 41 | 4L_TTA | caaaaacgattcctacccgtaatttttccatagttaatttc | 41 |
| 4L_CTA | gaaattaactatggaaaaactacgggtaggaatcgtttttg | 41 | 4L_TTG | caaaaacgattcctacccgcaatttttccatagttaatttc | 41 |
| 5R_CGA | ttaactatggaaaaactgcgagtaggaatcgtttttgg | 38 | 5R_CGA | ccaaaaacgattcctactcgcagtttttccatagttaa | 38 |
| 5R_AGG | ttaactatggaaaaactgagggtaggaatcgtttttgg | 38 | 5R_CGC | ccaaaaacgattcctacgcgcagtttttccatagttaa | 38 |
| 5R_AGA | attaactatggaaaaactgagagtaggaatcgtttttgg | 39 | 5R_CGT | caccaaaaacgattcctacacgcagtttttccatagttaat | 41 |
| 6V_GTG | ctatggaaaaactgcgggtgggaatcgtttttggtg | 36 | 6V_GTG | caccaaaaacgattcccacccgcagtttttccatag | 36 |
| 6V_GTC | ctatggaaaaactgcgggtcggaatcgtttttggtg | 36 | 6V_GTT | caccaaaaacgattccaacccgcagtttttccatag | 36 |
| 7G_GGG | ggaaaaactgcgggtagggatcgtttttggtggtaaatc | 39 | 7G_GGG | gatttaccaccaaaaacgatccctacccgcagtttttcc | 39 |
| 7G_GGC | ggaaaaactgcgggtaggcatcgtttttggtggtaaatc | 39 | 7G_GGT | gatttaccaccaaaaacgatacctacccgcagtttttcc | 39 |
| 8I_ATA | ggaaaaactgcgggtaggaatagtttttggtggtaaatcag | 41 | 8I_ATT | ctgatttaccaccaaaaacaattcctacccgcagtttttc | 40 |
| 9V_GTC | ctgcgggtaggaatcgtctttggtggtaaatcagc | 35 | 9V_GTC | gctgatttaccaccaaagacgattcctacccgcag | 35 |
| 9V_GTA | ctgcgggtaggaatcgtatttggtggtaaatcagc | 35 | 9V_GTG | gctgatttaccaccaaacacgattcctacccgcag | 35 |
| 10F_TTC | cgggtaggaatcgttttcggtggtaaatcagcg | 33 | 10F_TTC | cgctgatttaccaccgaaaacgattcctacccg | 33 |
| 11G_GGC | gggtaggaatcgtttttggcggtaaatcagcggaac | 36 | 11G_GGC | gttccgctgatttaccgccaaaaacgattcctaccc | 36 |
| 11G_GGA | gggtaggaatcgtttttggaggtaaatcagcggaac | 36 | 11G_GGG | gttccgctgatttacccccaaaaacgattcctaccc | 36 |
| 12G_GGC | gtaggaatcgtttttggtggcaaatcagcggaacatg | 37 | 12G_GGC | catgttccgctgatttgccaccaaaaacgattcctac | 37 |
| 12G_GGA | gtaggaatcgtttttggtggaaaatcagcggaacatg | 37 | 12G_GGG | catgttccgctgatttcccaccaaaaacgattcctac | 37 |
| 13K_AAG | gaatcgtttttggtggtaagtcagcggaacatgaag | 36 | 13K_AAG | cttcatgttccgctgacttaccaccaaaaacgattc | 36 |
| 14S_TCC | gaatcgtttttggtggtaaatccgcggaacatgaagtg | 38 | 14S_TCC | cacttcatgttccgcggatttaccaccaaaaacgattc | 38 |
| 14S_AGT | gaatcgtttttggtggtaaaagtgcggaacatgaagtg | 38 | 14S_TCG | cacttcatgttccgccgatttaccaccaaaaacgattc | 38 |
| 14S_AGC | gaatcgtttttggtggtaaaagcgcggaacatgaagtg | 38 | 14S_TCT | cacttcatgttccgcagatttaccaccaaaaacgattc | 38 |
| 15A_GCA | cgtttttggtggtaaatcagcagaacatgaagtgtctctgc | 41 | 15A_GCA | gcagagacacttcatgttctgctgatttaccaccaaaaacg | 41 |
| 15A_GCC | cgtttttggtggtaaatcagccgaacatgaagtgtctctg | 40 | 15A_GCT | gcagagacacttcatgttcagctgatttaccaccaaaaacg | 41 |
| 16E_GAG | ggtggtaaatcagcggagcatgaagtgtctctgc | 34 | 16E_GAG | gcagagacacttcatgctccgctgatttaccacc | 34 |
| 17H_CAC | gtggtaaatcagcggaacacgaagtgtctctgcaatc | 37 | 17H_CAC | gattgcagagacacttcgtgttccgctgatttaccac | 37 |
| 18E_GAG | ggtaaatcagcggaacatgaggtgtctctgcaatcg | 36 | 18E_GAG | cgattgcagagacacctcatgttccgctgatttacc | 36 |
| 19V_GTA | ggtaaatcagcggaacatgaagtatctctgcaatcggcaaaaaac | 45 | 19V_GTA | gtttttgccgattgcagagatacttcatgttccgctgatttacc | 45 |
| 19V_GTC | gtaaatcagcggaacatgaagtctctctgcaatcggcaaaaaac | 44 | 19V_GTT | gtttttgccgattgcagagaaacttcatgttccgctgatttac | 44 |
| 20S_AGC | gcggaacatgaagtgagcctgcaatcggcaaaaaac | 36 | 20S_AGC | gtttttgccgattgcaggctcacttcatgttccgc | 36 |
| 20S_AGT | gcggaacatgaagtgagtctgcaatcggcaaaaaac | 36 | 20S_TCA | gtttttgccgattgcagtgacacttcatgttccgc | 36 |

| | | | | | |
|---|---|---|---|---|---|
| 20S_TCC | gcggaacatgaagtgtccctgcaatcggcaaaaaac | 36 | S20S_TCG | gttttttgccgattgcagcgacacttcatgttccgc | 36 |
| 21L_CTA | cagcggaacatgaagtgtctctacaatcggcaaaaaacattg | 42 | 21L_CTA | caatgttttttgccgattgtagagacacttcatgttccgctg | 42 |
| 21L_CTC | cggaacatgaagtgtctctccaatcggcaaaaaacattgtc | 41 | 21L_CTT | caatgttttttgccgattgaagagacacttcatgttccgctg | 42 |
| 21L_TTA | cagcggaacatgaagtgtctttacaatcggcaaaaaacattgtc | 44 | 21L_TTG | gacaatgttttttgccgattgcaaagacacttcatgttccg | 41 |
| 22Q_CAG | ggaacatgaagtgtctctgcagtcggcaaaaaacattgtc | 40 | 22Q_CAG | gacaatgttttttgccgactgcagagacacttcatgttcc | 40 |
| 23S_AGC | catgaagtgtctctgcaaagcgcaaaaaacattgtcgatg | 40 | 23S_AGC | catcgacaatgttttttgcgctttgcagagacacttcatg | 40 |
| 23S_AGT | gaacatgaagtgtctctgcaaagtgcaaaaaacattgtcgatgc | 44 | 23S_TCA | gcatcgacaatgttttttgctgattgcagagacacttcatgttc | 44 |
| 23S_TCC | gaacatgaagtgtctctgcaatccgcaaaaaacattgtcgatg | 43 | S23S_TCT | gcatcgacaatgttttttgcagattgcagagacacttcatgttc | 44 |
| 24A_GCC | gtgtctctgcaatcggccaaaaacattgtcgatgc | 35 | 24A_GCC | gcatcgacaatgtttttggccgattgcagagacac | 35 |
| 24A_GCG | gtgtctctgcaatcggcgaaaaacattgtcgatgc | 35 | 24A_GCT | gcatcgacaatgtttttagccgattgcagagacacttcatg | 41 |
| 25K_AAG | gtctctgcaatcggcaaagaacattgtcgatgccattg | 38 | 25K_AAG | caatggcatcgacaatgttctttgccgattgcagagac | 38 |
| 26N_AAT | ctctgcaatcggcaaaaaatattgtcgatgccattgataaaag | 42 | 26N_AAT | cttttatcaatggcatcgacaatatttttgccgattgcagag | 42 |
| 27I_ATA | ctctgcaatcggcaaaaaacatagtcgatgccattgataaaag | 43 | 27I_ATC | cttttatcaatggcatcgacgatgttttttgccgattgcagag | 43 |
| 28V_GTA | ctgcaatcggcaaaaaacattgtagatgccattgataaaagtcgc | 45 | 28V_GTA | gcgacttttatcaatggcatctacaatgttttttgccgattgcag | 45 |
| 28V_GTG | caatcggcaaaaaacattgtggatgccattgataaaagtcgc | 42 | 28V_GTT | gcgacttttatcaatggcatcaacaatgttttttgccgattgcag | 45 |
| 29D_GAC | cggcaaaaaacattgtcgacgccattgataaaagtcgc | 38 | 29D_GAC | gcgacttttatcaatggcgtcgacaatgttttttgccg | 38 |
| 30A_GCA | cggcaaaaaacattgtcgatgcaattgataaaagtcgcttcg | 42 | 30A_GCA | cgaagcgacttttatcaattgcatcgacaatgttttttgccg | 42 |
| 30A_GCC | ggcaaaaaacattgtcgatgcgattgataaaagtcgcttcgac | 43 | 30A_GCT | gtcgaagcgacttttatcaatagcatcgacaatgttttttgccg | 44 |
| 31I_ATA | gcaaaaaacattgtcgatgccatagataaaagtcgcttcgacg | 43 | 31I_ATC | cgtcgaagcgacttttatcgatggcatcgacaatgttttttg | 42 |
| 32D_GAC | cattgtcgatgccattgacaaaagtcgcttcgacg | 35 | 32D_GAC | cgtcgaagcgacttttgtcaatggcatcgacaatg | 35 |
| 33K_AAG | cattgtcgatgccattgataagagtcgcttcgacgttg | 38 | 33K_AAG | caacgtcgaagcgactcttatcaatggcatcgacaatg | 38 |
| 34S_AGC | gtcgatgccattgataaaagccgcttcgacgttgtg | 36 | 34S_AGC | cacaacgtcgaagcggcttttatcaatggcatcgac | 36 |
| 34S_TCA | gtcgatgccattgataaatcacgcttcgacgttgtgc | 37 | 34S_TCC | cacaacgtcgaagcgggatttatcaatggcatcgac | 36 |
| 34S_TCG | gtcgatgccattgataaatcgcgcttcgacgttgtg | 36 | 34S_TCT | gcacaacgtcgaagcgagatttatcaatggcatcgac | 37 |
| 35R_AGA | gatgccattgataaaagtagattcgacgttgtgctgctgg | 40 | 35R_AGA | ccagcagcacaacgtcgaatctacttttatcaatggcatc | 40 |
| 35R_AGG | gtcgatgccattgataaaagtaggttcgacgttgtgctg | 39 | 35R_CGA | cagcacaacgtcgaatcgacttttatcaatggcatcgac | 39 |
| 35R_CGG | cgatgccattgataaaagtcggttcgacgttgtgctg | 37 | 35R_CGT | cagcacaacgtcgaaacgacttttatcaatggcatcgac | 39 |
| 36F_TTT | gccattgataaaagtcgctttgacgttgtgctgctg | 36 | 36F_TTT | cagcagcacaacgtcaaagcgacttttatcaatggc | 36 |
| 37D_GAT | ccattgataaaagtcgcttcgatgttgtgctgctggg | 37 | 37D_GAT | cccagcagcacaacatcgaagcgacttttatcaatgg | 37 |
| 38V_GTA | gataaaagtcgcttcgacgtagtgctgctgggcattg | 37 | 38V_GTA | caatgcccagcagcactacgtcgaagcgacttttatc | 37 |
| 38V_GTC | gataaaagtcgcttcgacgtcgtgctgctgggcattg | 37 | 38V_GTG | caatgcccagcagcaccacgtcgaagcgacttttatc | 37 |
| 39V_GTA | gataaaagtcgcttcgacgttgtactgctgggcattgataaac | 43 | 39V_GTA | gtttatcaatgcccagcagtacaacgtcgaagcgacttttatc | 43 |
| 39V_GTC | gataaaagtcgcttcgacgttgtcctgctgggcattgataaac | 43 | 39V_GTT | gtttatcaatgcccagcagaacaacgtcgaagcgacttttatc | 43 |
| 40L_CTA | gtcgcttcgacgttgtgctactgggcattgataaac | 36 | 40L_CTA | gtttatcaatgcccagtagcacaacgtcgaagcgac | 36 |
| 40L_CTC | cgcttcgacgttgtgctcctgggcattgataaac | 34 | 40L_CTT | gtttatcaatgcccagaagcacaacgtcgaagcg | 34 |
| 40L_TTA | gtcgcttcgacgttgtgttactgggcattgataaacaag | 39 | 40L_TTG | gtttatcaatgcccagcaacacaacgtcgaagcg | 34 |

| | | | | | |
|---|---|---|---|---|---|
| 41L_CTA | cttcgacgttgtgctgctaggcattgataaacaaggg | 37 | 41L_CTA | cccttgtttatcaatgcctagcagcacaacgtcgaag | 37 |
| 41L_CTC | cttcgacgttgtgctgctcggcattgataaacaagg | 36 | 41L_CTT | cccttgtttatcaatgccaagcagcacaacgtcgaag | 37 |
| 41L_TTA | cttcgacgttgtgctgttaggcattgataaacaagggc | 38 | 41L_TTG | cccttgtttatcaatgcccaacagcacaacgtcgaag | 37 |
| 42G_GGG | gacgttgtgctgctggggattgataaacaagggcaatg | 38 | 42G_GGG | cattgcccttgtttatcaatccccagcagcacaacgtc | 38 |
| 42G_GGA | gacgttgtgctgctgggaattgataaacaagggcaatg | 38 | 42G_GGT | cattgcccttgtttatcaatacccagcagcacaacgtc | 38 |
| 43I_ATA | gttgtgctgctgggcatagataaacaagggcaatg | 35 | 43I_ATC | cattgcccttgtttatcgatgcccagcagcacaac | 35 |
| 44D_GAC | gtgctgctgggcattgacaaacaagggcaatggc | 34 | 44D_GAC | gccattgcccttgtttgtcaatgcccagcagcac | 34 |
| 45K_AAG | ctgctgggcattgataagcaagggcaatggcac | 33 | 45K_AAG | gtgccattgcccttgcttatcaatgcccagcag | 33 |
| 46Q_CAG | gctgggcattgataaacaggggcaatggcacgtc | 34 | 46Q_CAG | gacgtgccattgcccctgtttatcaatgcccagc | 34 |
| 47G_GGC | ctgggcattgataaacaaggccaatggcacgtcagc | 36 | 47G_GGC | gctgacgtgccattggccttgtttatcaatgcccag | 36 |
| 47G_GGA | ctgggcattgataaacaaggacaatggcacgtcagc | 36 | 47G_GGT | gctgacgtgccattgaccttgtttatcaatgcccag | 36 |
| 48Q_CAG | gcattgataaacaagggcagtggcacgtcagcgatg | 36 | 48Q_CAG | catcgctgacgtgccactgcccttgtttatcaatgc | 36 |
| 50H_CAT | gataaacaagggcaatggcatgtcagcgatgccag | 35 | 50H_CAT | ctggcatcgctgacatgccattgcccttgtttatc | 35 |
| 51V_GTG | gataaacaagggcaatggcacgtgagcgatgccagcaattatc | 43 | 51V_GTG | gataattgctggcatcgctcacgtgccattgcccttgtttatc | 43 |
| 51V_GTA | caagggcaatggcacgtaagcgatgccagcaattatc | 37 | 51V_GTT | gataattgctggcatcgctaacgtgccattgcccttg | 37 |
| 52S_TCC | ggcaatggcacgtctccgatgccagcaattatc | 33 | 52S_TCC | gataattgctggcatcggagacgtgccattgcc | 33 |
| 52S_TCA | ggcaatggcacgtctcagatgccagcaattatc | 33 | 52S_TCG | gataattgctggcatccgagacgtgccattgcc | 33 |
| 52S_AGT | ggcaatggcacgtcagtgatgccagcaattatc | 33 | 52S_TCT | gataattgctggcatcagagacgtgccattgcc | 33 |
| 53D_GAC | caatggcacgtcagcgacgccagcaattatctgc | 34 | 53D_GAC | gcagataattgctggcgtcgctgacgtgccattg | 34 |
| 54A_GCG | ggcacgtcagcgatgcgagcaattatctgctaaatg | 36 | 54A_GCG | catttagcagataattgctcgcatcgctgacgtgcc | 36 |
| 54A_GCA | ggcacgtcagcgatgcaagcaattatctgctaaatg | 36 | 54A_GCT | catttagcagataattgctagcatcgctgacgtgcc | 36 |
| 55S_TCC | cacgtcagcgatgcctccaattatctgctaaatgcag | 37 | 55S_TCC | ctgcatttagcagataattggaggcatcgctgacgtg | 37 |
| 55S_TCA | cacgtcagcgatgcctcaaattatctgctaaatgcag | 37 | 55S_TCG | ctgcatttagcagataattcgaggcatcgctgacgtg | 37 |
| 55S_AGT | cacgtcagcgatgccagtaattatctgctaaatgcag | 37 | 55S_TCT | ctgcatttagcagataattagaggcatcgctgacgtg | 37 |
| 56N_AAC | gtcagcgatgccagcaactatctgctaaatgcagac | 36 | 56N_AAC | gtctgcatttagcagatagttgctggcatcgctgac | 36 |
| 57Y_TAC | cagcgatgccagcaattacctgctaaatgcagacg | 35 | 57Y_TAC | cgtctgcatttagcaggtaattgctggcatcgctg | 35 |
| 58L_CTT | cgatgccagcaattatcttctaaatgcagacgatcctg | 38 | 58L_CTT | caggatcgtctgcatttagaagataattgctggcatcg | 38 |
| 58L_CTC | cgatgccagcaattatctcctaaatgcagacgatcc | 36 | 58L_TTA | ggatcgtctgcatttagtaaataattgctggcatcgc | 37 |
| 58L_CTA | cgatgccagcaattatctactaaatgcagacgatcctg | 38 | 58L_TTG | ggatcgtctgcatttagcaaataattgctggcatcg | 36 |
| 59L_CTT | gatgccagcaattatctgcttaatgcagacgatcctg | 37 | 59L_CTT | caggatcgtctgcattaagcagataattgctggcatc | 37 |
| 59L_CTG | gatgccagcaattatctgctgaatgcagacgatcctg | 37 | 59L_TTA | caggatcgtctgcatttaacagataattgctggcatcg | 38 |
| 59L_CTC | gatgccagcaattatctgctcaatgcagacgatcctg | 37 | 59L_TTG | caggatcgtctgcattcaacagataattgctggcatc | 37 |
| 60N_AAC | ccagcaattatctgctaaacgcagacgatcctgc | 34 | 60N_AAC | gcaggatcgtctgcgtttagcagataattgctgg | 34 |
| 61A_GCG | cagcaattatctgctaaatgcggacgatcctgcccatattg | 41 | 61A_GCG | caatatgggcaggatcgtccgcatttagcagataattgctg | 41 |
| 61A_GCC | cagcaattatctgctaaatgccgacgatcctgcccatattg | 41 | 61A_GCT | caatatgggcaggatcgtcagcatttagcagataattgctg | 41 |
| 62D_GAT | caattatctgctaaatgcagatgatcctgcccatattgcg | 40 | 62D_GAT | cgcaatatgggcaggatcatctgcatttagcagataattg | 40 |

| | | | | | |
|---|---|---|---|---|---|
| 64P_CCC | gctaaatgcagacgatcccgcccatattgcgttg | 34 | 64P_CCC | caacgcaatatgggcgggatcgtctgcatttagc | 34 |
| 64P_CCA | gctaaatgcagacgatccagcccatattgcgttg | 34 | 64P_CCG | caacgcaatatgggccggatcgtctgcatttagc | 34 |
| 65A_GCG | ctaaatgcagacgatcctgcgcatattgcgttgcgc | 36 | 65A_GCG | gcgcaacgcaatatgcgcaggatcgtctgcatttag | 36 |
| 65A_GCA | ctaaatgcagacgatcctgcacatattgcgttgcgc | 36 | 65A_GCT | gcgcaacgcaatatgagcaggatcgtctgcatttag | 36 |
| 66H_CAC | cagacgatcctgcccacattgcgttgcgccc | 31 | 66H_CAC | gggcgcaacgcaatgtgggcaggatcgtctg | 31 |
| 123A_GCG | gaatgctgcgggtcgcgaatttaccgtttgtagg | 34 | 123A_GCG | cctacaaacggtaaattcgcgacccgcagcattc | 34 |
| 123A_GCA | gaatgctgcgggtcgcaaatttaccgtttgtagg | 34 | 123A_GCT | cctacaaacggtaaattagcgacccgcagcattc | 34 |
| 124N_AAC | gctgcgggtcgccaacttaccgtttgtaggttc | 33 | 124N_AAC | gaacctacaaacggtaagttggcgacccgcagc | 33 |
| 125L_CTG | ctgcgggtcgccaatctgccgtttgtaggttctg | 34 | 125L_CTG | cagaacctacaaacggcagattggcgacccgcag | 34 |
| 125L_CTC | ctgcgggtcgccaatctcccgtttgtaggttctg | 34 | 125L_CTT | cagaacctacaaacggaagattggcgacccgcag | 34 |
| 125L_CTA | ctgcgggtcgccaatctaccgtttgtaggttctg | 34 | 125L_TTG | cagaacctacaaacggcaaattggcgacccgcag | 34 |
| 126P_CCC | cgggtcgccaatttaccctttgtaggttctgatgttc | 37 | 126P_CCC | gaacatcagaacctacaaagggtaaattggcgacccg | 37 |
| 126P_CCA | cgggtcgccaatttaccatttgtaggttctgatgttc | 37 | 126P_CCT | gaacatcagaacctacaaaaggtaaattggcgacccg | 37 |
| 127F_TTC | gtcgccaatttaccgttcgtaggttctgatgttctg | 36 | 127F_TTC | cagaacatcagaacctacgaacggtaaattggcgac | 36 |
| 128V_GTG | cgccaatttaccgtttgtgggttctgatgttctgg | 35 | 128V_GTG | ccagaacatcagaacccacaaacggtaaattggcg | 35 |
| 128V_GTC | cgccaatttaccgtttgtcggttctgatgttctgg | 35 | 128V_GTT | ccagaacatcagaaccaacaaacggtaaattggcg | 35 |
| 129G_GGC | ccaatttaccgtttgtaggctctgatgttctggcttc | 37 | 129G_GGC | gaagccagaacatcagagcctacaaacggtaaattgg | 37 |
| 129G_GGA | ccaatttaccgtttgtaggatctgatgttctggcttcag | 39 | 129G_GGG | gaagccagaacatcagaccctacaaacggtaaattgg | 37 |
| 130S_TCA | ccaatttaccgtttgtaggttcagatgttctggcttcag | 39 | 130S_TCA | ctgaagccagaacatctgaacctacaaacggtaaattgg | 39 |
| 130S_AGT | ccaatttaccgtttgtaggtagtgatgttctggcttcag | 39 | 130S_TCC | ctgaagccagaacatcggaacctacaaacggtaaattg | 38 |
| 130S_AGC | caatttaccgtttgtaggtagcgatgttctggcttcag | 38 | 130S_TCG | ctgaagccagaacatccgaacctacaaacggtaaattg | 38 |
| 132V_GTC | cgtttgtaggttctgatgtcctggcttcagcagc | 34 | 132V_GTC | gctgctgaagccaggacatcagaacctacaaacg | 34 |
| 132V_GTA | cgtttgtaggttctgatgtactggcttcagcagc | 34 | 132V_GTG | gctgctgaagccagcacatcagaacctacaaacg | 34 |
| 133L_CTT | gtttgtaggttctgatgttcttgcttcagcagcctg | 36 | 133L_CTT | caggctgctgaagcaagaacatcagaacctacaaac | 36 |
| 133L_CTC | gtttgtaggttctgatgttctcgcttcagcagcctg | 36 | 133L_TTA | catacaggctgctgaagctaaaacatcagaacctacaaac | 40 |
| 133L_CTA | gtttgtaggttctgatgttctagcttcagcagcctg | 36 | 133L_TTG | caggctgctgaagccaaaacatcagaacctacaaac | 36 |
| 134A_GCC | gtaggttctgatgttctggcctcagcagcctgtatg | 36 | 134A_GCC | catacaggctgctgaggccagaacatcagaacctac | 36 |
| 134A_GCA | gtaggttctgatgttctggcatcagcagcctgtatg | 36 | 134A_GCG | catacaggctgctgacgccagaacatcagaacctac | 36 |
| 135S_TCC | gttctgatgttctggcttccgcagcctgtatggac | 35 | 135S_TCC | gtccatacaggctgcggaagccagaacatcagaac | 35 |
| 135S_AGT | gttctgatgttctggctagtgcagcctgtatggac | 35 | 135S_TCG | gtccatacaggctgccgaagccagaacatcagaac | 35 |
| 135S_AGC | gttctgatgttctggctagcgcagcctgtatggac | 35 | 135S_TCT | gtccatacaggctgcagaagccagaacatcagaac | 35 |
| 136A_GCG | ctgatgttctggcttcagcggcctgtatggacaaag | 36 | 136A_GCG | ctttgtccatacaggccgctgaagccagaacatcag | 36 |
| 136A_GCC | ctgatgttctggcttcagccgcctgtatggacaaag | 36 | 136A_GCT | ctttgtccatacaggcagctgaagccagaacatcag | 36 |
| 137A_GCG | gttctggcttcagcagcgtgtatggacaaagatgtc | 36 | 137A_GCG | gacatctttgtccatacacgctgctgaagccagaac | 36 |
| 137A_GCA | gttctggcttcagcagcatgtatggacaaagatgtc | 36 | 137A_GCT | gacatctttgtccatacaagctgctgaagccagaac | 36 |
| 138C_TGC | ctggcttcagcagcctgcatggacaaagatgtcac | 35 | 138C_TGC | gtgacatctttgtccatgcaggctgctgaagccag | 35 |

| | | | | | |
|---|---|---|---|---|---|
| 140D_GAT | cttcagcagcctgtatggataaagatgtcaccaaacg | 37 | 140D_GAT | cgtttggtgacatctttatccatacaggctgctgaag | 37 |
| 141K_AAG | gcagcctgtatggacaaggatgtcaccaaacgtc | 34 | 141K_AAG | gacgtttggtgacatccttgtccatacaggctgc | 34 |
| 142D_GAC | cagcctgtatggacaaagacgtcaccaaacgtctg | 35 | 142D_GAC | cagacgtttggtgacgtctttgtccatacaggctg | 35 |
